# Dynamics of Take-off in Bipedal Animals and Robots

**DOI:** 10.64898/2026.05.07.723416

**Authors:** Guan-Yu Chen, Zhi-Yong Wu, Shan-Hung Chen, Patricia J. Yang

**Affiliations:** Department of Power Mechanical Engineering, National Tsing Hua University, Taiwan

## Abstract

Take-off is a fast and energy-efficient strategy for bipedal animals, such as birds, to achieve rapid movement; however, how muscle physiology scales to govern this universal behavior remains unresolved. Research in other species’ physiologies is not readily applicable. As a result, important questions about whether theropod dinosaurs, such as *Tyrannosaurus rex*, were capable of jumping remain unanswered. In this article, we coupled Lagrangian dynamics with Hill’s muscle equations. We developed new experimental methods to quantify joint rotational stiffness and damping, thereby enabling a systematic description of lower-limb mechanics. The approach establishes a novel kinetic framework that links muscle contractile properties to lower-limb performance without invoking control optimization. Animal observations and tabletop mechanisms verify the framework. The mechanics model reveals that the take-off time of about 0.1 s across body masses of 0.003 to 90 kg is achievable, as heavier birds generate proportionally higher reaction forces. Additionally, *Tyrannosaurus rex* should be capable of jumping, based on the available physiology data. Beyond evolutionary insights, our framework provides a new methodology for analyzing the mechanical properties of biological joints and informing the design of scalable bio-inspired robots.

## 1. Introduction

Jumping is a fundamental locomotor strategy used by bipedal animals for rapid locomotion, predator evasion, and accessing elevated substrates. Effective take-off requires the rapid conversion of muscular work into whole-body kinetic energy, integrating muscle dynamics, joint mechanics, and body inertia across diverse morphologies. Understanding these interactions is critical not only for biomechanics but also for bio-inspired robotics^1,2^, assistive devices^3^, and the interpretation of locomotor performance in extinct species.

Previous studies have elucidated key mechanisms that shape jump performance at the muscle and tendon levels. Joint mechanical advantage constrains how muscle forces are transmitted to the ground, influencing whether movement relies primarily on direct muscle actuation or elastic energy storage^4^. Experimental work in frogs has shown that intrinsic muscle properties alone do not fully determine performance, highlighting the importance of system-level interactions^5^. Tendon elasticity modulates muscle operating lengths, enabling favorable force–length and force–velocity function^6^, and direct evidence of elastic energy storage demonstrates catapult-like mechanisms in vertebrates^7^. Broader syntheses emphasize the diverse functional roles of biological springs in vertebrate locomotion^8^. At cross-scales, Sutton et al. showed that latch-mediated spring-actuated systems benefit small animals but become energetically limiting in larger ones^9^, whereas Bobbert demonstrated that force–velocity constraints alone predict declining jump performance at small sizes, underscoring the importance of size-dependent muscle dynamics^10^. Importantly, these studies treat muscle physiology as a limiting factor on performance, but do not embed it as a constitutive constraint within the equations governing take-off dynamics. Similarly, previous experimental work has quantified tendon^11,12^ or single-joint mechanics^13,14^. While valuable, these studies measure local tissue properties in isolation and cannot directly predict whole-body take-off dynamics across body sizes. This limitation highlights the need for a framework that integrates local mechanical measurements with system-level dynamics, allowing for predictive, cross-scale modeling of take-off performance.

Recent advances in predictive musculoskeletal modeling have enabled cross-species simulations of locomotion using high-dimensional, muscle-driven frameworks. Using OpenSim-based models, Clemente et al. employed predictive musculoskeletal simulations to reveal mechanistic links among posture, energetic cost, and running speed across extant mammals^15^. Extending these approaches to jumping, Bishop et al. developed a muscle-driven model of a generalized ground-dwelling bird (tinamou) to evaluate feasible jumping strategies and joint coordination patterns^16^. Their model used optimal control, with sensitivity analyses of selected muscle properties, but represented only one generalized bird and did not address body-size effects on take-off dynamics.

Haldane et al. showed with a series-elastic jumping robot that elastic storage and changes in mechanical advantage can redistribute motor work over time and amplify peak output power^17^. Polet and Labonte used an idealized constant-gearing model to analyze how muscle force, shortening speed, work capacity, and external loading affect the optimal transmission ratio in robotic galago^18^. These studies provide mechanism-level insights into elasticity and gearing rather than cross-species predictions of take-off dynamics.

The need for such a cross-scale framework is especially apparent in studies of extinct dinosaurs, for which movement and soft-tissue properties cannot be measured directly. Sellers et al. used a species-specific forward-dynamics model and optimized muscle activation patterns to reconstruct slow locomotion in the giant sauropod *Argentinosaurus*^19^. In the absence of passive support structures, the 83-tonne model required restricted joint ranges of motion to generate a feasible gait. Cuff et al. subsequently reviewed experimental and computational studies of archosaur locomotion and identified predictive musculoskeletal simulation as a promising approach for investigating walking, running, and jumping in extinct taxa^20^. They also emphasized the need to apply these approaches to a broader range of archosaur taxa and body forms. More recently, Charles et al. applied optimal-control simulation directly to vertical jumping in the approximately 13 kg early theropod *Coelophysis bauri*^21^. The predicted performance was comparable to that of a generalized extant bird and was sensitive to proximal tail-joint range of motion and relative tail mass. Together, these studies establish the value of species-specific simulations for evaluating locomotor capabilities in extinct animals, but do not provide a common framework for cross-scale comparison.

Here, we investigate whether common scaling patterns of bipedal take-off emerge across Aves, Macropodidae, and Primates, which span a broad range of body sizes and lower-limb proportions. We first quantify rotational stiffness and damping at the toe, ankle, and knee across species. We then incorporate these measurements into a reduced-order model that couples Hill-type muscle with constrained Lagrangian dynamics. The model is evaluated using cross-species high-speed observations and tabletop mechanisms. By integrating empirical joint mechanics, muscle dynamics, morphological scaling, and ground-contact constraints, the framework enables interpretable comparisons across morphologically diverse taxa and body sizes and supports its application to extinct theropods.

## 2. Results

### 2.1 Lower-limb compound pendulum model

We employ Lagrangian dynamics to simulate variations in joint angles during take-off, aiming to clarify the influence of Hill’s equation^22^ on lower-limb function and to capture the trajectory of limb movements throughout the process (Fig. 1B). To reduce complexity, we restrict the model to sagittal-plane motion and impose bilateral symmetry^23^, allowing the animal to be represented by a half-model. The upper body is represented by a point mass fixed relative to the thigh, with no independent upper-body rotational degree of freedom. The position of the upper-body center of mass, defined by *D* and *ϕ*, is determined from static equilibrium and the geometry of the initial squat posture, as detailed in Supplementary Materials. The lower limb is simplified as a compound pendulum with five generalized coordinates: the orientations of the three segments relative to the horizontal axis (*θ*_1_, *θ*_2_, *θ*_3_) and the horizontal and vertical positions of the toe (*x*_*toe*_, *y*_*toe*_). During ground contact, both toe coordinates are constrained while the three segments remain free to rotate. The corresponding Lagrange multipliers represent the horizontal and vertical components of the ground reaction force^24^. When the vertical ground reaction force falls below the prescribed toe-off threshold^25^, the contact constraints are removed, and all five generalized coordinates become dynamic during the aerial phase. Because soft tissues (e.g., tendons, ligaments, cartilage) around joints provide resistance and store elastic energy during rotation^5,8,26^, we incorporate rotational stiffness *k*_*t*_ and damping *c*_*t*_ at each joint to better approximate realistic joint behavior. The simplified model is illustrated in Fig. 1A.

### 2.2 Cross-species joint mechanics

In the simulations, *k*_*t*_ and *c*_*t*_ are implemented as equivalent passive properties, whereas active muscle force is calculated separately using the Hill force–velocity relationship. No series-elastic tendon element was explicitly modelled; therefore, the passive effects of the retained tendons entered the model only through the measured joint properties. To characterize the mechanical properties of joints across species, we measured passive rotational stiffness *k*_*t*_ and damping *c*_*t*_ of the knee, ankle, and toe. (Supplementary Tables 6-7). Rotational stiffness is determined from torque–angle relations during externally imposed passive rotation, consistent with the rotational analog of Hooke’s law τ = *k*_*t*_ Δ*θ* One measurement of the knee joint is shown as an example (Fig. 2A). Figs. 2C-E show the relationships between torque and angular displacement of the toe, ankle, and knee joints, respectively. The slope of the fitted line based on these data defines the joint’s rotational stiffness.

The measured *k*_*t*_ represents the equivalent passive rotational stiffness of the joint as a whole, rather than the stiffness of an individual tissue. Because the muscles were not activated during testing, the torque–angle relationship reflects the combined passive contributions of muscle–tendon units, ligaments, joint capsules, and other periarticular tissues. Using Hooke’s law and the definition of Young’s modulus, we derive an idealized relationship linking joint rotational stiffness to equivalent anatomical structure (see Section 4.3):

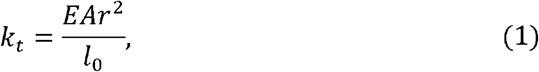

where *E, A*, and *l*_*0*_ are Young’s modulus, cross-sectional area, and original length of the soft tissues. Assuming that animals are isometric, the predicted rotational stiffness of the joint *k*_*t*_ ~***M***^***1***^ The experimental value of rotational stiffness at the knee provides the scaling relationship *k*_*t3*_ ~***M*** ^1.09^. The exponent 1.09 is nearly unity, indicating isometry. However, we found that the rotational stiffness of the toe provides the scaling relationship *k*_*t1*_ ~***M***^0.52^, and the rotational stiffness of the ankle provides *k*_*t2*_ ~***M***^0.63^. These values are not close to unity; therefore, *k*_*t2*_ and *k*_*t3*_ are not isometric. These results may be attributed to the characteristics of the anatomical structure. As observed in anatomical samples, Avian lower limbs exhibit significant sectional differences in soft tissue distribution. The thigh and shank regions are densely packed with muscles, tendons, and ligaments, while the tarsometatarsus and toes are characterized by elongated structures and lightweight adaptations with relatively sparse muscles, tendons, and ligaments. Consequently, a greater amount of soft tissue is engaged and stretched during rotational motion at the knee joint. The elastic properties of these soft tissues contribute to the higher rotational stiffness at the knee joint, facilitating the storage of elastic potential energy during locomotion.

Rotational damping coefficients were extracted from free-vibration decay using logarithmic decrement analysis (see Section 4.3). As shown in Fig. 2B, we mount the thigh on the stand, adjusting the fixed angle to allow the shank to hang and swing freely. Fig. 2F-H show the amplitude decrement with time of the toe, ankle, and knee joint free vibration, respectively. The experimental values of the rotational damping coefficient at toe *c*_*t1*_ ~*M*^1.23^, at ankle *c*_*t2*_ ~*M*^***1*.*57***^, and at knee *c*_*t*_ ~***M***^1.44^. Since damping ratio ζ is a dimensionless parameter ranging from 0 to 1, *c*_*t*_ is proportional to *c*_*ct*_. Assuming isometry (*k*_*t*_ ~*M*^1^, moment of inertia ~***M***^5/3^), the predicted 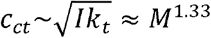. The derived exponent demonstrates close agreement with the experimental results.

### 2.3 Cross-scale animal observation

We collected take-off time *T*, maximum ground reaction force *F*, and maximum velocity *V*_max_ across bipedal species mainly in Aves, Macropodidae, and Primates. These three focal groups represent distinct lower-limb morphologies and span more than four orders of magnitude in body mass. For take-off time, we filmed the take-off of 27 species of animals using high-speed videography at zoos (Supplementary Videos 4-6), obtained 9 data points from the literature, and 2 videos from YouTube. For maximum ground reaction force, we obtained 11 data points from the literature and 13 data points for maximum velocity. All raw data are listed in Supplementary Tables 2–4. The data we collected included multiple species, and therefore, we applied the phylogenetic generalized least squares (PGLS) to the dataset. PGLS accounts for the similarities between species that arise from their shared ancestry, adjusting the regression model’s slope and intercept, providing more accurate estimates of trait relationships^24^.

The observational dataset includes jumps with or without a preparatory countermovement. For the cross-species scaling analysis, take-off is defined from the onset of upward body motion to toe-off, excluding any preceding downward countermovement from the measured take-off time (Fig. 3A). Figs. 3B-D show the results of the cross-scale animal observations, together with the corresponding scaling exponents *b* and their 95% confidence intervals, indicating significant positive scaling with body mass. For the take-off time data, the smallest animal is a hummingbird with 0.003 kg, and the largest is a human with 90 kg. The take-off time ranges from 28.3 ms to 295 ms, and the average is 121.2 ± 49.7 ms (mean ± S.D., n=38). Using the PGLS method, we derive the scaling function *T* = 128.5 *M* ^0.18^(ms), *F* = 40.4 *M* ^1.02^(N), and *V*_max_ = 2.6*M*^0.15^ (m/s). Maximum ground reaction force scales approximately in direct proportion to body mass, as the confidence interval for its exponent includes unity, whereas take-off time and maximum velocity increase more weakly with body mass. This weak mass dependence is also evident in the raw observations: despite a nearly 30,000-fold variation in body mass, take-off time varies by only approximately tenfold (28.3–295 ms), with more than 50% of the observations falling between 101 and 150 ms. The observed scaling relationships were also mechanically consistent with approximation using the impulse–momentum relationship. During the take-off process, the hindlimbs exert a force on the ground. This force equals the ground reaction force, and according to our definition of the take-off phase, the velocity change is *V*_max_. Therefore, take-off time can be approximated as:

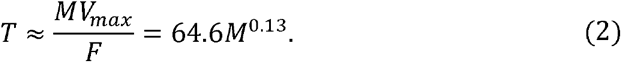

The trend of Eq. 2 closely matches the animal observation result but is approximately half of the value overall (Fig. 3B). This difference arises because Eq. 2 uses the reported maximum ground reaction force, whereas the exact impulse–momentum relationship depends on the average net force acting throughout the take-off phase. Because the peak force is greater than the average net force, Eq. 2 underestimates the absolute take-off time. Nevertheless, it captures the observed scaling trend.

### 2.4 Cross-scale simulations

After deriving the equations of motion (see Section 4.2), we solve the joint dynamics numerically to obtain lower-limb trajectories during take-off. All simulations represent squat jumps (SJs), which begin from a static crouched posture. In contrast, countermovement jumps (CMJs) include a preparatory downward movement before propulsion^28,29^. CMJs achieve greater jump heights because muscle activation and force develop before shortening begins. However, the advantage over SJs is modest and does not produce an order-of-magnitude difference in performance^28^. This study focuses on general mechanical patterns across body mass. We therefore do not consider the additional effects of countermovement.

We set up the posture as a controlled initial condition (see Section 4.4). Within each taxonomic group, all body masses use the same initial angles (*θ*_1_, *θ*_2_, *θ*_3_). The model therefore does not incorporate body-size-dependent changes in initial posture or joint range of motion. Larger animals may adopt more extended limb postures, which increase effective mechanical advantage but reduce the available distance for limb extension and mechanical work^15,30^. This standardization allowed body-mass-dependent morphological, joint, and muscle scaling relationships to vary while keeping other factors constant.

The initial segment angles for Aves are obtained from published guinea-fowl kinematics^23^. Those for Primates are estimated from a representative frame in publicly available squat-jump footage^31^. Comparable postural data are unavailable for Macropodidae, so the model uses the avian initial posture. All muscles are activated simultaneously in each simulation to provide a common basis for comparison, even though animals alter the sequence of muscle contractions in response to different situations^32,33^. All model parameters are scaled with body mass using the functions in Supplementary Tables 8–10. Aves, Macropodidae, and Primates use distinct lower-limb scaling functions to account for differences in limb proportions.

The model generates take-off time, maximum velocity during take-off, and maximum ground reaction force for body masses corresponding to 38 species (Supplementary Table 11). The fitting equations, scaling exponents, 95% confidence intervals, and sample sizes are reported in Table 1. All simulation trials take-off sucessfully without falling or swinging. The model describes the coordination between muscle contraction and lower-limb movement during take-off. We then compare the simulations with animal observations (Fig. 4). The take-off videos in simulations are shown in Supplementary Videos 1–3.

**Table 1.**
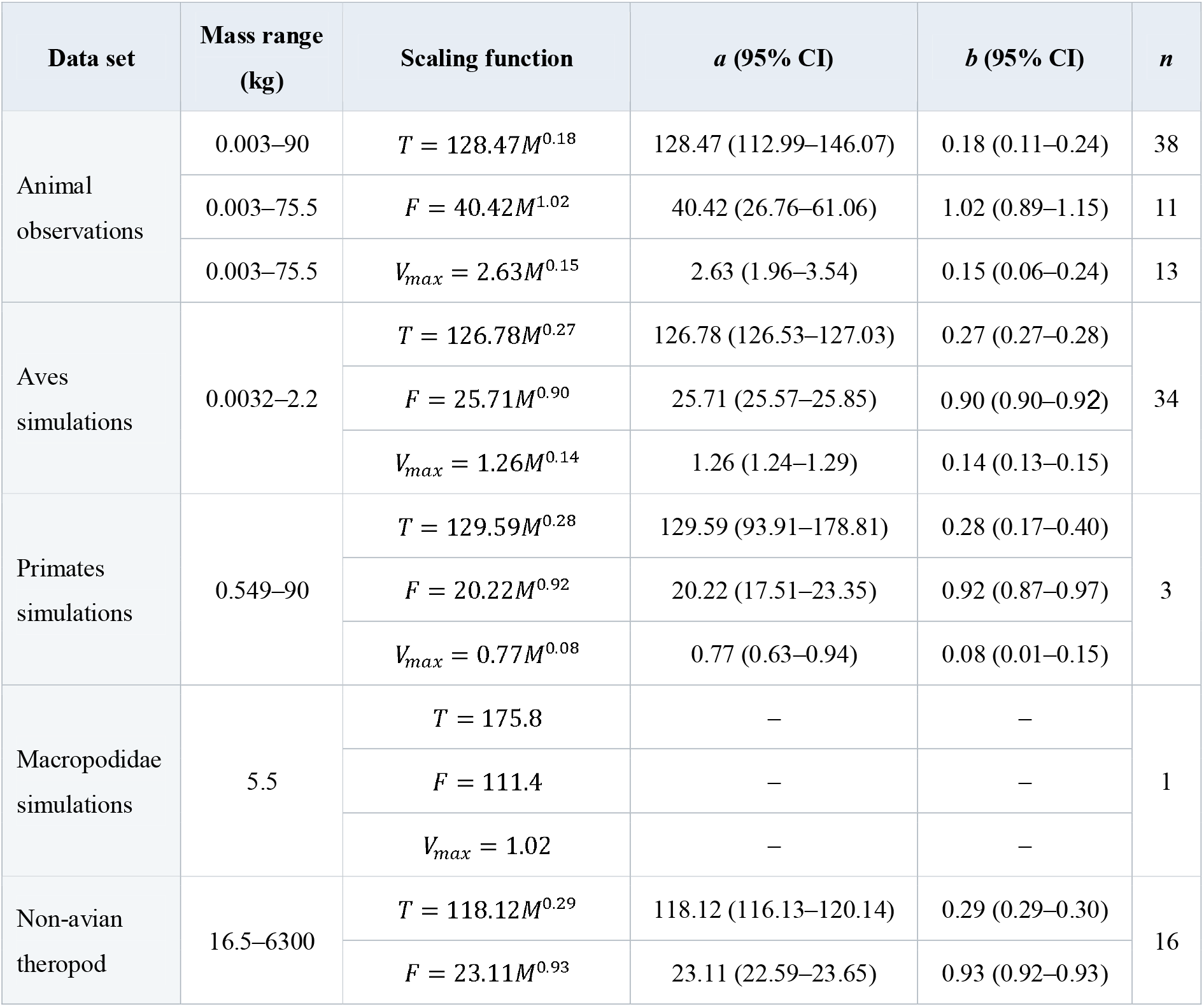
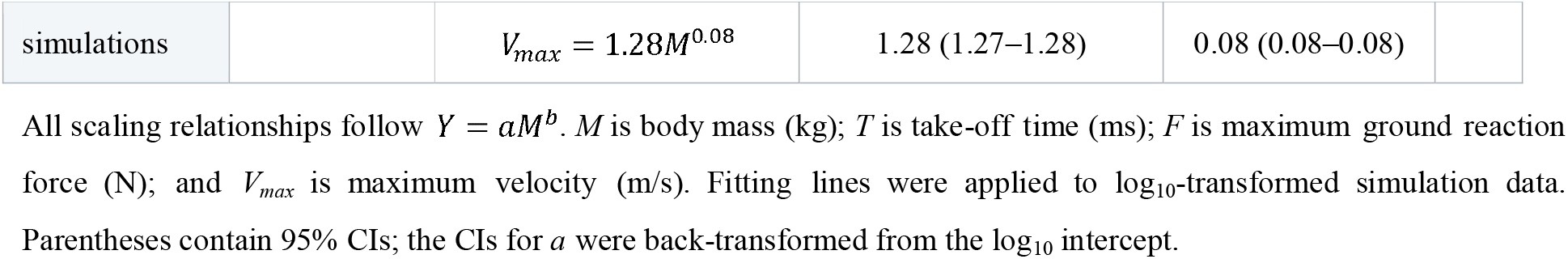
Scaling relationships for observed and simulated take-off performance.

Despite this overall agreement, the correspondence between observed and simulated scaling relationships varies across take-off variables. For take-off time, the observations, Aves simulation, and Primates simulation fits have similar prefactors of 128.47, 126.78, and 129.59, respectively, indicating close agreement in magnitude. However, the simulated exponents for Aves and Primates are 0.27 and 0.28, respectively, both exceeding the observed exponent of 0.18 and its 95% confidence interval of 0.11–0.24. The Aves confidence interval (0.27–0.28) does not overlap the observed interval, whereas the Primates interval (0.17–0.40) does. For maximum ground reaction force, the Aves and Primates exponents of 0.90 and 0.92 both fall within the 95% confidence interval of the observed exponent of 1.02 (0.89–1.15). The model thus reproduces a near-proportional scaling of maximum ground reaction force with body mass, although the lower simulated prefactors indicated that absolute force magnitudes were generally underestimated. Maximum velocity shows similar agreement in its scaling trend. The simulated Aves exponent of 0.14 is nearly identical to the observed exponent of 0.15, while the Primates exponent of 0.08 also falls within the observed 95% confidence interval of 0.06–0.24. The three observed primate data points in Fig. 4 show a similarly shallow trend to the Primates simulation, qualitatively supporting the predicted within-group scaling. However, the lower simulated relationships indicate that maximum velocities are underestimated. Accordingly, *V*_max_/ (*Tg*) decreases only slightly with body mass in the observations but declines more strongly in the simulations, reflecting the combined effects of steeper take-off-time scaling and lower simulated velocities.

Discrepancies in maximum ground reaction force may arise because the simulations assign the same initial posture to all species within each group, although actual posture varies among species and individuals in reality. Such variation affects effective mechanical advantage and the range of joint extension. Differences in maximum velocity may arise because the model fixes the upper body relative to the thigh, limiting the contribution of lower-limb propulsion to whole-body acceleration. Overall, the model reproduces the cross-scale trends in maximum ground reaction force and velocity, as well as the magnitude of take-off time. The remaining discrepancies reflect both model limitations and the expected differences between controlled simulations and reality.

### 2.5 Extension to non-avian theropod body masses

We further extend the model to non-avian theropods based on comparisons with extant animals. Aves are the only extant theropod lineage, and non-avian theropods and Aves share similarities in hindlimb skeletal architecture and in the scaling of skeletal morphology with body mass^34^. Previous studies have inferred theropod locomotion from fossil limb mechanics and comparisons with modern birds. Farlow et al. showed that smaller theropods were generally more agile, whereas larger taxa faced greater mechanical constraints^35^. Avian-based predictive models have also incorporated body size and speed to infer the locomotor capabilities of extinct theropods^36^. However, locomotor, speed, and limb loading alone cannot determine whether take-off can be completed. Take-off also depends on the dynamic interaction between muscle contraction and lower-limb movement. To address this gap, we use published body-mass estimates for 16 non-avian theropods, ranging from 16.5 to 6,300 kg in the literature^37^, as body mass *M* in the model. The morphological, joint, and muscle parameters are assigned using the Aves scaling relationships summarized in Supplementary Tables 8–10. These relationships, including their reported coefficients and exponents, were treated as prescribed model inputs rather than extrapolations of the Aves output-fitting lines. We then evaluated whether each mass-specific simulation met the prescribed take-off criterion as shown in Section 4.4. The simulation results are listed in Supplementary Table 11.

The simulations for Aves and non-avian theropods show similar scaling patterns in take-off time and maximum ground reaction force. The corresponding scaling exponents are 0.27 and 0.29 for take-off time and 0.90 and 0.93 for maximum ground reaction force, respectively (Table 1). Maximum velocity increases more slowly with body mass in non-avian theropods (*M*^0.08^) than in Aves (*M*^0.14^). Maximum ground reaction force (*M*^0.9^) increases slightly less than body mass (*M*^*1*^) in both groups. Similarly, the dimensionless acceleration *V*_max_/ (*Tg*) decreases as *M*^−0.13^ in the Aves simulations and as *M*^−0.21^ in the non-avian theropod simulations. Thus, as body mass increases, the simulated theropods generate greater forces over longer take-off periods, but their acceleration progressively declines, as shown in Fig. 4D.

All 16 non-avian theropod models complete take-off. The 6,300 kg *Tyrannosaurus rex* model completes take-off at 1.56 s, with a maximum ground reaction force of 79.69 kN and a maximum velocity of 2.54 m/s. Based on the published body-mass estimate and the avian-derived assumptions for morphology, joint mechanics, and muscle properties, the theropods in simulations are capable of take-off.

This conclusion depends on the model inputs. The output relationships for non-avian theropods were fitted directly to simulations in body mass of 16.5–6,300 kg, whereas the avian-derived input relationships used to construct the models extended beyond the body-mass range of the extant animals from which they were derived. The 95% confidence intervals of the output-fitting lines reflect uncertainty in the estimated regression trends of the simulation outputs; they do not represent the probability that an extinct theropod could or could not jump. These results should therefore be interpreted as conditional predictions under a specific set of avian-derived scaling assumptions, rather than evidence that *Tyrannosaurus rex* could jump under every biologically plausible combination of morphological and muscle parameters.

### 2.6 Tabletop verification

To evaluate the feasibility of the compound pendulum model under physical systems and enable its application to bioinspired robotics or engineering systems across multiple scales, we make 3 tabletop mechanisms differing in body mass by an order of magnitude and compare their take-off times with animals of corresponding mass (Supplementary Fig. 8). We simplify the compound pendulum model and replicate each segment according to the scaling functions and the Aves skeleton. Each mechanism is subjected to three repeated trials, and the take-off times are shown in Supplementary Table 12. The experimental results closely align with the data from animal observation, as shown in Fig. 4A. The take-off times of bipedal animals are related to the dimensions and proportions of their body features. Larger animals bear heavier loads but are also capable of generating greater force, so the increase in take-off time with body mass is relatively slow.

## 3. Conclusions

This study provides a cross-scale mechanical explanation of bipedal take-off. Across 38 species spanning nearly five orders of magnitude in body mass, take-off time remains within the same order of magnitude, while maximum ground reaction force increases approximately in proportion to body mass. The passive rotational stiffness and damping scale with body mass. The take-off performance emerges from the dynamic interaction among body-size-dependent morphology, active muscle contraction, passive joint mechanics, and body inertia, rather than from body mass or muscle force alone.

We integrate the experimentally measured joint stiffness and damping, Hill-type muscle dynamics, morphological scaling, and ground-contact constraints within a reduced-order forward-dynamics model. Without re-optimizing movement coordination for each body mass, the model reproduces the observed scaling patterns in maximum ground reaction force and maximum velocity, as well as take-off-time magnitude. However, it predicted a stronger mass dependence of take-off time and lower absolute force and velocity than observed. Tabletop mechanisms provided additional physical support by producing take-off times comparable to animal observations.

Under published body-mass estimates and the prescribed avian-derived scaling relationships, all 16 non-avian theropod models completed take-off, including the 6,300 kg *Tyrannosaurus rex* model. This result describes the predicted performance of an avian-scaled musculoskeletal model and may change if the actual morphology or muscle properties of *Tyrannosaurus rex* differed from these assumptions. Overall, this framework links local muscle and joint mechanics to whole-body take-off and provides an interpretable basis for investigating locomotion in extinct animals and designing bio-inspired mechanisms across scales.

## 4. Methods

### 4.1 Animal observation

This study filmed 27 data points on bipedal animals’ take-off time at the Taipei Zoo in Taiwan and at Zoo Atlanta in the US. We filmed one video per individual. A high-speed camera (Phantom VEO-E310L), lens (Canon Macro 100 mm), and tripod were used to capture videos of birds taking off and recording the corresponding timings at 1000 frames per second. All procedures were approved by the IACUC of National Tsing Hua University (Approval Number: 111027-1) and the Georgia Institute of Technology (Protocol Number: A16017).

### 4.2 Compound pendulum model

We construct a simplified dynamic model in MATLAB (R2023a, MathWorks). Joint rotational stiffness, damping, and segment masses of Aves were measured in Section 4.3 (Supplementary Tables 5–7), whereas muscle contractile parameters, limb dimensions, and segment masses of primates and macropodidae were obtained from published morphological datasets. All scaling functions and corresponding sources are summarized in Supplementary Tables 8–10.

The configuration of the model was described by five generalized coordinates,

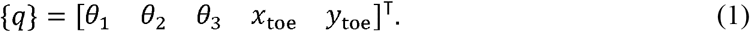

Here, *q*_*i*_ is the th generalized coordinate. *θ*_1_, *θ*_2_, and *θ*_3_ are the angles between the tarsus, shank, and thigh and the horizontal axis, respectively; *x*_*toe*_ and *y*_*toe*_ are the horizontal and vertical positions of the toe. The translational and angular velocities of each segment were obtained by differentiating their positions with respect to time.

During ground contact, the toe is fixed at the origin; therefore, its horizontal and vertical coordinates satisfy

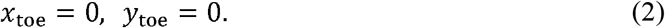

These two coordinates are retained in the generalized-coordinate vector so that the same five-coordinate formulation could be used during both ground contact and the aerial phase. They were held fixed only during ground contact and became unconstrained after toe-off. To incorporate the two contact conditions into the equations of motion, we defined the constraint Jacobian

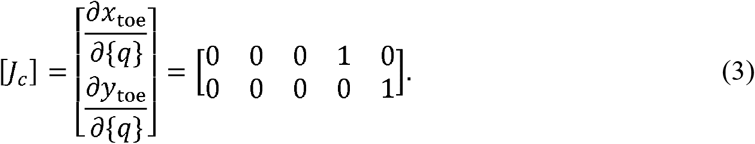

In the present model, [*J*_*c*_] is simply a coordinate-selection matrix: the first row selects the horizontal toe coordinate, and the second row selects the vertical toe coordinate. The two reaction forces to maintain the toe at the origin were represented by the Lagrange multipliers *λ*_*x*_and *λ*_*y*_, collected in {*λ*} = [*λ*_*x*_ *λ*_*y*_]^T^. The transpose [*J*_*c*_]^T^ maps these two contact forces onto the five generalized-coordinate directions. In this formulation, it places *λ*_*x*_and *λ*_*y*_in the equations governing *x*_toe_and *y*_toe_, respectively. The constrained Lagrange equations can therefore be written as

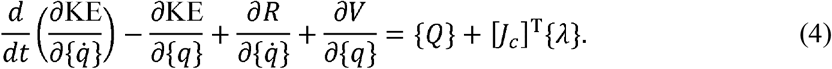

Here, *KE* is the kinetic energy, *V* is the potential energy, and is Rayleigh’s dissipation function^38^. is the vector of nonconservative generalized forces, specified below from the muscle-generated torques. The kinetic energy, potential energy, and dissipation function were

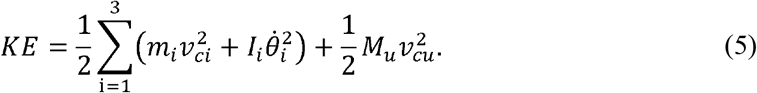

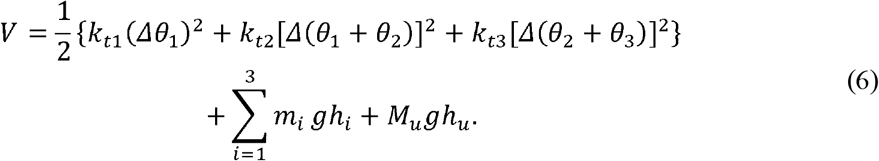

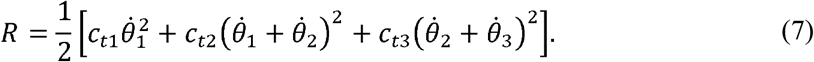

Here, *m*_*i*_, *l*_*i*_, *v*_*ci*_, *I*_*i*_, and *h*_*i*_ are the mass, length, center-of-mass velocity, moment of inertia about the center of mass, and center-of-mass height of segment, respectively; 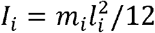. Similarly, *M*_*u*_, *v*_*cu*_, and *h*_*u*_ are the mass, center-of-mass velocity, and center-of-mass height of the upper-body point mass. The quantities *k*_*ti*_ and *c*_*ti*_ are the rotational stiffness and damping coefficients, respectively. Substituting Eqs. (5) – (7) into Eq. (4) and reorganizing the equations of motion:

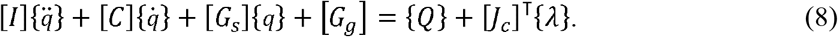

Here, is the inertia matrix, is the Coriolis, centrifugal, and damping matrix, [*G*_*s*_] is the stiffness matrix, and [*G*_*g*_] is the gravitational matrix. The term [*J*_*c*_]^⊤^{*λ*} is the generalized force associated with the ground-contact constraints. is constant, so that when we differentiate the position constraints twice gives zero horizontal and vertical toe accelerations during ground contact. The model has initial conditions 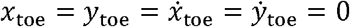. At each evaluation during numerical integration, the five equations of motion were combined with the two acceleration constraints and solved as

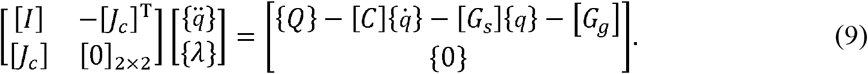

The five equations of motion and two constraint equations are solved simultaneously for the five generalized accelerations and two Lagrange multipliers. Because the constraints are imposed directly on the horizontal and vertical positions of the toe, the Lagrange multipliers correspond to the horizontal and vertical components of the ground reaction force,

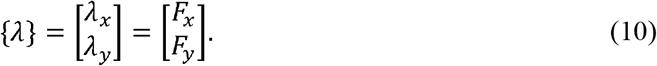

The ground reaction force is obtained directly from the constrained equations of motion rather than reconstructed separately from segment accelerations. This interpretation of the contact multipliers follows Oshima et al.^24^. When the toe-off criterion is satisfied (see Section 4.4), the two contact constraints and their associated multipliers are removed. The generalized coordinates and velocities at toe-off are retained as the initial conditions for the aerial phase, during which the equations of motion become

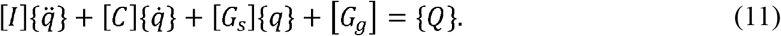

Muscle contraction generates torques only in the three rotational coordinates,

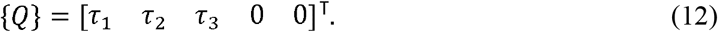

The torques exerted during joint rotation come from muscle contraction. According to the hindlimb anatomy, the muscles primarily responsible for extending the hindlimb are divided into two groups: those extending the knee joint and those extending the ankle joint. The free-body diagram in Supplementary Fig. 1 illustrates the torque exerted on each joint. The force from the thigh, *F*_3_, and the force from the calf, *F*_2_, generate torques on both the proximal and distal parts. The equations are shown as follows:

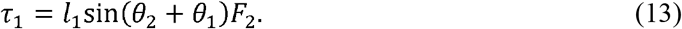

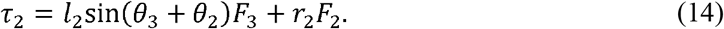

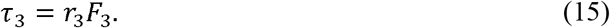

Here, *r*_2_ is the radius of the ankle joint, and *r*_3_ is the radius of the knee joint. Because they are approximately half the epicondylar length, we assume the joint radius is approximately 1.5 times the bone shaft radius^39^.

The model requires three components to satisfy the initial state and muscle-generated forces. First, static equilibrium at the prescribed squat posture is used to determine the position of the upper-body point mass, described by *D* and *ϕ*, together with the muscle forces required to maintain the initial posture, *F*_2,static_ and *F*_3,static_ Second, the Hill force-velocity relationship^22^ is reformulated for implementation in the compound pendulum model by relating muscle shortening velocity to segment rotation and converting muscle tension to force using muscle cross-sectional area. Third, previous experimental studies of muscle activation are reviewed to formulate a time-dependent activation function for take-off^9,26^, whose initial value is determined from the muscle forces required for static equilibrium. Together, these components provide the muscle forces and corresponding generalized torques required by the equations of motion. Their detailed formulations are presented in Supplementary Information.

### 4.3 Measurements of mechanical properties of joints

We measure the rotational stiffness *k*_*t*_ and rotational damping *c*_*t*_ by experiments. The *k*_*t*_ and *c*_*t*_ experiments incorporate three kinds of joints: knee, ankle, and toe. For each joint, we perform individual experiments. For each experiment, we have 5 to 8 specimens, and these specimens are from the following species: chicken (*Gallus gallus domesticus*) from supermarket; cattle egret (*Bubulcus coromandus*) and sacred ibis (*Threskiornis aethiopicus*) from the College of Life Science and Medicine at NTHU; muller’s barbet (*Psilopogon nuchalis*), gray treepie (*Dendrocitta formosae*), black bulbul (*Hypsipetes leucocephalus*), Malaysian night heron (*Gorsachius melanolophus*), and spotted dove (*Spilopelia chinensis*) from Hsinchu Wild Bird Federation. The masses of birds range from 0.06 kg to 3 kg. The results are provided in Supplementary Tables 6 and 7. Chen et al. found that one to two freeze-thaw cycles have minimal impact on soft tissues, while repeated cycles can damage the soft tissues’ microstructure^41^. Wren expected that at room temperature, the mechanical properties of biological tissues exhibit minimal differences compared to those in vivo^42^. Wang found that the moduli of wallaby and tiger tail tendons remained constant for temperatures from 20°C to 41°C^43^. In this study, all samples were stored at –18°C and thawed at room temperature in zip-seal bags before experiments to prevent desiccation. Experiments were conducted at room temperature, and samples were discarded after each test.

To measure *k*_*t*_ of a specific lower-limb joint, the segment proximal to the joint is securely fixed to a supporting frame, while the joint is allowed to rotate freely in the plane of measurement. The limb portion distal to the joint was fixed so that its constituent segments move together as a single unit. For example, when measuring knee-joint stiffness, the thigh is fixed to the supporting frame, while the shank and the remaining distal portion of the limb are fixed together so that motion occurs only at the knee. The distal portion of the limb moves through the required range of motion without contacting the supporting frame or any surrounding objects. We attached a rope to the distal portion of the limb at one end, and suspended weights at the other end. The applied load is gradually increased, and the corresponding change in joint angle is recorded. When the joint rotates, the surrounding soft tissues are stretched, with the elongation being the product of the bone radius *r* and the angular displacement Δ*θ*. The stretching force is the soft tissues’ spring constant multiplied by the elongation: *f*= *kr*Δ*θ*. Therefore, when a torque τ is applied to the joint, the torque can be expressed not only as the rotational stiffness multiplied by the angular displacement, *kt*Δ θ, but also as the spring force,*krΔ θ*, multiplied by the joint’s radius, which is approximately equal to the bone radius *r*. The formula is written as τ=*k*_*t*_Δ*θ*=*kr*^2^ Δ*θ* According to Hooke’s law and the definition of Young’s modulus, 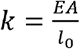 where *E* is Young’s modulus, *A* is the cross-sectional area, and *l*_0_ is the original length of the soft tissues. Finally, we derive the relationship between the rotational stiffness of a joint and its anatomical structure:

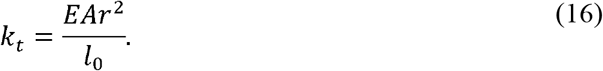

Assuming that animals are isometric, *A*~*M*^2/3^, *r*~*M*^1/3^ and *l*_0_ ~*M*^1/3^. *E* is constant across animal size. Thus, the predicted rotational stiffness of the joint *kt*~*M*^1^

We derive the rotational damping coefficient of joints by utilizing the Logarithmic decrement δ, which represents the rate at which the amplitude of a free-damped vibration decreases. The function is written as

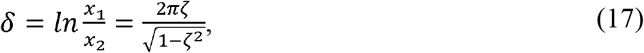

where *x*_1_and *x*_2_ are the amplitudes of two adjacent free vibrations, respectively, and the damping ratio 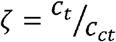. The critical rotational damping constant 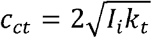 where *I*_i_ is the moment of inertia of the rotational component, which is 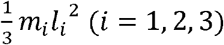.We mount the thigh on the stand, adjusting the fixed angle to allow the shank to hang and swing freely. Then we raise and release the shank, allowing it to swing naturally. The amplitude of its swing gradually diminished due to damping. The amplitudes of the first two peaks of each vibration are substituted into Eq. 27 to determine the damping ratio ζ and then obtain the rotational damping coefficient *c*_*t*_ of each joint.

### 4.4 Simulation across scales

Simulations were performed across body masses for Aves, Macropodidae, and Primates to examine general patterns of squat-jump dynamics. Each simulation required initial values and velocities of the generalized coordinates {*q*^(0)^}=[*θ*_1,0_ *θ*_2,0_ *θ*_3,0_*x*_*toe,o*_ *y*_*toe,0*_]^T^,Whole-animal body mass and selection of the taxonomic group. All initial velocities were zero for squat jumps. Taxonomic group affected only the scaling relationships of lower-limb segment masses and skeletal lengths. All remaining morphological, joint, and muscle parameters were estimated from body mass using the same scaling functions across the three groups (Supplementary Tables 8–10). Avian and primate postures were based on guinea-fowl kinematics^23^ and human squat-jump footage^31^, respectively. Because comparable cross-scale macropodid data were unavailable, macropodids use the avian posture. These prescribed postures serve only as controlled initial conditions and do not imply that extant animals use a fixed take-off posture.

During ground contact, toe coordinates are constrained, and the Lagrange multipliers provide the horizontal and vertical ground reaction forces. When *F*_*y*_ falls below one-thousandth of the model weight, calculated using half of the body mass, both constraints are removed, and the model switches to unconstrained motion, allowing the toe to move freely. Take-off is recorded only when exceeds 10^−6^, rather than zero as the numerical threshold.

The simulation ends if the ankle angle *θ*_1_ + *θ*_2_ or knee angle *θ*_2_+ *θ*_3_ reaches 180^°^, representing the physiological limit of joint extension. After take-off, the simulation ends when *y*_toe_ reaches half of the metatarsal length. Each simulation provided the time histories of joint angles, toe coordinates, ground reaction forces, and upper-body point-mass velocity. Slow-motion animation is also generated for inspection of the simulated movement, as shown in Supplementary video 1–3.

### 4.5 Tabletop mechanism

In addition to impulse-momentum theory and numerical simulation, we constructed cross-scale tabletop mechanisms and measured their take-off times. Each mechanism represents one side of a bilaterally symmetric bird: one hindlimb and half of the upper-body mass. The target animal body masses of 0.05, 0.5, and 5 kg were selected to span two orders of magnitude rather than to represent three particular bird species. Accordingly, the total mass of each mechanism was set to half of the animal body mass, resulting in mechanism masses of 0.025, 0.25, and 2.5 kg, respectively. Because the thigh is the shortest of the three lower-limb segments and experiences the smallest torque, we simplified the three-segment compound-pendulum model by removing the thigh. The segment masses and spring forces of the mechanisms were determined from the scaling relationships and the Aves parameters provided in Supplementary Tables 8–10. The lower-limb components were fabricated by 3D printing with carbon-fiber material, springs were used to mimic muscle forces, and a paper-clay mass block was mounted on the limb to represent the upper-body mass. A string connecting the ends of the two lower-limb bars held the mechanism in its initial configuration by counteracting the spring force. Cutting the string released the mechanism, allowing the spring force to generate torque about the ankle joint and initiate take-off. Each mechanism was tested in three repeated trials, and the measured take-off times are reported in Supplementary Table 12.

## 5. Data availability

All data are presented in the paper, Supplementary Figures, and Supplementary Tables.

## 6. Code availability

The code used in this study is available on Zenodo at https://doi.org/10.5281/zenodo.22282038

## Acknowledgment

We thank Zoo Atlanta and Taipei Zoo for providing the environment and samples for animal filming. Thanks to Hsinchu Wild Bird Federation for providing biological specimens for the measurements of mechanical properties of joints. Thanks to Chen Siang Ng for providing biological specimens and suggestions. Thanks to David Hu for providing photography equipment to film in Zoo Atlanta. Thanks to Catherine Grey for her initial help with data collection in Zoo Atlanta. Thanks to Yum Ji Chan, KuoAn Wu, and Feifei Qian for their valuable feedback and suggestions. Thanks to my lab mates for their help with data collection in Taipei Zoo and suggestions.

## References

1. Shin WD, Phan H-V, Daley MA, Ijspeert AJ, Floreano D. 2024 Fast ground-to-air transition with avian-inspired multifunctional legs. Nature 636, 86–91. (doi:10.1038/s41586-024-08228-9).

2. Ramirez Serrano F, Hyun N-sP, Steinhardt E, Lechère P-L, Wood RJ. 2025 A springtail-inspired multimodal walking-jumping microrobot. Sci. Robot. 10, eadp7854. (doi:10.1126/scirobotics.adp7854).

3. Andrade RM, Bonato P. 2021 The role played by mass, friction, and inertia on the driving torques of lower-limb gait training exoskeletons. IEEE Trans. Med. Robot. Bionics 3, 125–136. (doi:10.1109/TMRB.2021.3052014).

4. Olberding JP, Deban SM, Rosario MV, Azizi E. 2019 Modeling the determinants of mechanical advantage during jumping: consequences for spring- and muscle-driven movement. Integr. Comp. Biol. 59, 1515–1524. (doi:10.1093/icb/icz139).

5. Roberts TJ, Abbott EM, Azizi E. 2011 The weak link: do muscle properties determine locomotor performance in frogs? Phil. Trans. R. Soc. B 366, 1488–1495. (doi:10.1098/rstb.2010.0326).

6. Azizi E, Roberts TJ. 2010 Muscle performance during frog jumping: influence of elasticity on muscle operating lengths. Proc. R. Soc. B 277, 1523–1530. (doi:10.1098/rspb.2009.2051).

7. Astley HC, Roberts TJ. 2012 Evidence for a vertebrate catapult: elastic energy storage in the plantaris tendon during frog jumping. Biol. Lett. 8, 386–389. (doi:10.1098/rsbl.2011.0982).

8. Roberts TJ, Azizi E. 2011 Flexible mechanisms: the diverse roles of biological springs in vertebrate movement. J. Exp. Biol. 214, 353–361. (doi:10.1242/jeb.038588).

9. Sutton GP, Mendoza E, Azizi E, Longo SJ, Olberding JP, Ilton M, Patek SN. 2019 Why do large animals never actuate their jumps with latch-mediated springs? Because they can jump higher without them. Integr. Comp. Biol. 59, 1609–1618. (doi:10.1093/icb/icz145).

10. Bobbert MF. 2013 Effects of isometric scaling on vertical jumping performance. PLoS One 8, e71209. (doi:10.1371/journal.pone.0071209).

11. Matson A, Konow N, Miller S, Konow PP, Roberts TJ. 2012 Tendon material properties vary and are interdependent among turkey hindlimb muscles. J. Exp. Biol. 215, 3552–3558. (doi:10.1242/jeb.072728).

12. Vereecke EE, Channon AJ. 2013 The role of hind limb tendons in gibbon locomotion: springs or strings? J. Exp. Biol. 216, 3971–3980. (doi:10.1242/jeb.083527).

13. Schmitz RJ, Ficklin TK, Shimokochi Y, Nguyen A-D, Beynnon BD, Perrin DH, Shultz SJ. 2008 Varus/valgus and internal/external torsional knee joint stiffness differs between sexes. Am. J. Sports Med. 36, 1380–1388. (doi:10.1177/0363546508317411).

14. Liu C-L, Zhou J-P, Sun P-T, Chen B-Z, Zhang J, Tang C-Z, Zhang Z-J. 2020 Influence of different knee and ankle ranges of motion on the elasticity of triceps surae muscles, Achilles tendon, and plantar fascia. Sci. Rep. 10, 6643. (doi:10.1038/s41598-020-63730-0).

15. Clemente CJ, De Groote F, Dick TJ. 2024 Predictive musculoskeletal simulations reveal the mechanistic link between speed, posture and energetics among extant mammals. Nat. Commun. 15, 8594. (doi:10.1038/s41467-024-52924-z).

16. Bishop PJ, Falisse A, De Groote F, Hutchinson JR. 2021 Predictive simulations of musculoskeletal function and jumping performance in a generalized bird. Integr. Org. Biol. 3, obab006. (doi:10.1093/iob/obab006).

17. Haldane DW, Plecnik MM, Yim JK, Fearing RS. 2016 Robotic vertical jumping agility via series-elastic power modulation. Sci. Robot. 1, eaag2048. (doi:10.1126/scirobotics.aag2048).

18. Polet DT, Labonte D. 2024 Optimal gearing of musculoskeletal systems. Integr. Comp. Biol. 64, 987–1006. (doi:10.1093/icb/icae072).

19. Sellers WI, Margetts L, Coria RA, Manning PL. 2013 March of the titans: the locomotor capabilities of sauropod dinosaurs. PLoS One 8, e78733. (doi:10.1371/journal.pone.0078733).

20. Cuff AR, Demuth OE, Michel K, Otero A, Pintore R, Polet DT, Wiseman ALA, Hutchinson JR. 2022 Walking—and running and jumping—with dinosaurs and their cousins, viewed through the lens of evolutionary biomechanics. Integr. Comp. Biol. 62, 1281–1305. (doi:10.1093/icb/icac049).

21. Charles JP, Polet DT, Hutchinson JR. 2026 Form–function relationships determining optimal jumping performance in an early bipedal dinosaur. J. R. Soc. Interface 23, 20250918. (doi:10.1098/rsif.2025.0918).

22. Cohen C, Mouterde T, Quéré D, Clanet C. 2015 Capillary muscle. Proc. Natl Acad. Sci. USA 112, 6301–6306. (doi:10.1073/pnas.1419315112).

23. Henry HT, Ellerby DJ, Marsh RL. 2005 Performance of guinea fowl Numida meleagris during jumping requires storage and release of elastic energy. J. Exp. Biol. 208, 3293–3302. (doi:10.1242/jeb.01764).

24. Oshima T, Toriumi K, Fujikawa T, Momose N. 2004 Effects of the lower leg bi-articular muscle in jumping. J. Robot. Mechatron. 16, 643–648. (doi:10.20965/jrm.2004.p0643).

25. Koch MW, Leyendecker S. 2013 Structure preserving simulation of monopedal jumping. Arch. Mech. Eng. 60, 127–146. (doi:10.2478/meceng-2013-0008).

26. Rosario MV, Sutton GP, Patek SN, Sawicki GS. 2016 Muscle-spring dynamics in time-limited, elastic movements. Proc. R. Soc. B 283, 20161561. (doi:10.1098/rspb.2016.1561).

27. Cox SM, Easton KL, Lear MC, Marsh RL, Delp SL, Rubenson J. 2019 The interaction of compliance and activation on the force-length operating range and force generating capacity of skeletal muscle: a computational study using a guinea fowl musculoskeletal model. Integr. Org. Biol. 1, obz022. (doi:10.1093/iob/obz022).

28. Bobbert MF, Gerritsen KGM, Litjens MCA, van Soest AJ. 1996 Why is countermovement jump height greater than squat jump height? Med. Sci. Sports Exerc. 28, 1402–1412. (doi:10.1097/00005768-199611000-00009).

29. Barker LA, Harry JR, Mercer JA. 2018 Relationships between countermovement jump ground reaction forces and jump height, reactive strength index, and jump time. J. Strength Cond. Res. 32, 248–254. (doi:10.1519/JSC.0000000000002160).

30. Hutchinson JR. 2004 Biomechanical modeling and sensitivity analysis of bipedal running ability. I. Extant taxa. J. Morphol. 262, 421–440. (doi:10.1002/jmor.10241).

31. LocomotorLabTCU. 2012 Slow motion video of vertical jump with synchronized vertical force data. YouTube. See https://www.youtube.com/watch?v=qN3apht8zRs (accessed 4 September 2026).

32. Zajac FE. 1993 Muscle coordination of movement: a perspective. J. Biomech. 26 (Suppl. 1), 109–124. (doi:10.1016/0021-9290(93)90083-Q).

33. Bobbert MF, van Ingen Schenau GJ. 1988 Coordination in vertical jumping. J. Biomech. 21, 249–262. (doi:10.1016/0021-9290(88)90175-3).

34. Christiansen P. 1999 Long bone scaling and limb posture in non-avian theropods: evidence for differential allometry. J. Vertebr. Paleontol. 19, 666–680. (doi:10.1080/02724634.1999.10011180).

35. Farlow JO, Gatesy SM, Holtz Jr TR, Hutchinson JR, Robinson JM. 2000 Theropod locomotion. Am. Zool. 40, 640–663. (doi:10.1093/icb/40.4.640).

36. Bishop PJ, Graham DF, Lamas LP, Hutchinson JR, Rubenson J, Hancock JA, Wilson RS, Hocknull SA, Barrett RS, Lloyd DG et al. 2018 The influence of speed and size on avian terrestrial locomotor biomechanics: predicting locomotion in extinct theropod dinosaurs. PLoS One 13, e0192172. (doi:10.1371/journal.pone.0192172).

37. Christiansen P, Fariña RA. 2004 Mass prediction in theropod dinosaurs. Hist. Biol. 16, 85–92. (doi:10.1080/08912960412331284313).

38. Minguzzi E. 2015 Rayleigh’s dissipation function at work. Eur. J. Phys. 36, 035014. (doi:10.1088/0143-0807/36/3/035014).

39. Lee JH, Kim YS, Jeong YG, Lee NS, Han SY, Tubbs RS, Han SH. 2014 Sex determination from partial segments and maximum femur lengths in Koreans using computed tomography. Folia Morphol. 73, 353–358. (doi:10.5603/FM.2014.0052).

40. Hill AV. 1938 The heat of shortening and the dynamic constants of muscle. Proc. R. Soc. Lond. B 126, 136–195. (doi:10.1098/rspb.1938.0050).

41. Chen L, Wu Y, Yu J, Jiao Z, Ao Y, Yu C, Wang J, Cui G. 2011 Effect of repeated freezing–thawing on the Achilles tendon of rabbits. Knee Surg. Sports Traumatol. Arthrosc. 19, 1028–1034. (doi:10.1007/s00167-010-1278-y).

42. Wren TAL, Yerby SA, Beaupré GS, Carter DR. 2001 Mechanical properties of the human Achilles tendon. Clin. Biomech. 16, 245–251. (doi:10.1016/S0268-0033(00)00089-9).

43. Wang XT, de Ruister MR, Alexander RMcN, Ker RF. 1991 The effect of temperature on the tensile stiffness of mammalian tail tendons. J. Zool. 223, 491–497. (doi:10.1111/j.1469-7998.1991.tb04779.x).

44. Tobalske BW, Altshuler DL, Powers DR. 2004 Take-off mechanics in hummingbirds (Trochilidae). J. Exp. Biol. 207, 1345–1352. (doi:10.1242/jeb.00889).

45. Provini P, Tobalske BW, Crandell KE, Abourachid A. 2012 Transition from leg to wing forces during take-off in birds. J. Exp. Biol. 215, 4115–4124. (doi:10.1242/jeb.074484).

46. Earls KD. 2000 Kinematics and mechanics of ground take-off in the starling Sturnis vulgaris and the quail Coturnix coturnix. J. Exp. Biol. 203, 725–739. (doi:10.1242/jeb.203.4.725).

47. Boulinguez-Ambroise G, Boyer DM, Dunham NT, Yapuncich GS, Bradley-Cronkwright M, Zeininger A, Schmitt D, Young JW. 2024 Biomechanical and morphological determinants of maximal jumping performance in callitrichine monkeys. J. Exp. Biol. 227, jeb247413. (doi:10.1242/jeb.247413).

48. McGowan CP, Baudinette RV, Usherwood J, Biewener AA. 2005 The mechanics of jumping versus steady hopping in yellow-footed rock wallabies. J. Exp. Biol. 208, 2741–2751. (doi:10.1242/jeb.01702).

49. Demes B, Jungers WL, Fleagle JG, Wunderlich RE, Richmond BG. 1996 Body size and leaping kinematics in Malagasy vertical clingers and leapers. J. Hum. Evol. 31, 367–388. (doi:10.1006/jhev.1996.0066).

50. Scholz MN, D’Août K, Bobbert MF, Aerts P. 2006 Vertical jumping performance of bonobo (Pan paniscus) suggests superior muscle properties. Proc. R. Soc. B 273, 2177–2184. (doi:10.1098/rspb.2006.3568).

51. Heglund NC, Cavagna GA, Taylor CR. 1982 Energetics and mechanics of terrestrial locomotion. III. Energy changes of the centre of mass as a function of speed and body size in birds and mammals. J. Exp. Biol. 97, 41–56. (doi:10.1242/jeb.97.1.41).

52. Askew GN, Marsh RL, Ellington CP. 2001 The mechanical power output of the flight muscles of blue-breasted quail (Coturnix chinensis) during take-off. J. Exp. Biol. 204, 3601–3619. (doi:10.1242/jeb.204.21.3601).

53. Williams EV, Swaddle JP. 2003 Moult, flight performance and wingbeat kinematics during take-off in European starlings Sturnus vulgaris. J. Avian Biol. 34, 371–378. (doi:10.1111/j.0908-8857.2003.02964.x).

54. Nelson FE, Gabaldón AM, Roberts TJ. 2004 Force–velocity properties of two avian hindlimb muscles. Comp. Biochem. Physiol. A Mol. Integr. Physiol. 137, 711–721. (doi:10.1016/j.cbpb.2004.02.004).

55. Reiser PJ, Greaser ML, Moss RL. 1996 Contractile properties and protein isoforms of single fibres from the chicken pectoralis red strip muscle. J. Physiol. 493, 553–562. (doi:10.1113/jphysiol.1996.sp021403).

56. Marsh RL, Bennett AF. 1985 Thermal dependence of isotonic contractile properties of skeletal muscle and sprint performance of the lizard Dipsosaurus dorsalis. J. Comp. Physiol. B 155, 541–551. (doi:10.1007/BF00694443).

57. Askew GN, Marsh RL. 1997 The effects of length trajectory on the mechanical power output of mouse skeletal muscles. J. Exp. Biol. 200, 3119–3131. (doi:10.1242/jeb.200.24.3119).

58. Ranatunga KW. 1998 Temperature dependence of mechanical power output in mammalian (rat) skeletal muscle. Exp. Physiol. 83, 371–376. (doi:10.1113/expphysiol.1998.sp004120).

59. Wells JB. 1965 Comparison of mechanical properties between slow and fast mammalian muscles. J. Physiol. 178, 252–269. (doi:10.1113/jphysiol.1965.sp007626).

60. Close RI. 1969 Dynamic properties of fast and slow skeletal muscles of the rat after nerve cross-union. J. Physiol. 204, 331–346. (doi:10.1113/jphysiol.1969.sp008916).

61. Luff AR. 1981 Dynamic properties of the inferior rectus, extensor digitorum longus, diaphragm and soleus muscles of the mouse. J. Physiol. 313, 161–171. (doi:10.1113/jphysiol.1981.sp013656).

62. Luff AR. 1975 Dynamic properties of fast and slow skeletal muscles in the cat and rat following cross-reinnervation. J. Physiol. 248, 83–96. (doi:10.1113/jphysiol.1975.sp010963).

63. Buller AJ, Kean CJC, Ranatunga KW. 1987 Transformation of contraction speed in muscle following cross-reinnervation; dependence on muscle size. J. Muscle Res. Cell Motil. 8, 504–516. (doi:10.1007/BF01567909).

64. Jarvis JC. 1993 Power production and working capacity of rabbit tibialis anterior muscles after chronic electrical stimulation at 10 Hz. J. Physiol. 470, 157–169. (doi:10.1113/jphysiol.1993.sp019852).

65. Sréter FA, Luff AR, Gergely J. 1975 Effect of cross-reinnervation on physiological parameters and on properties of myosin and sarcoplasmic reticulum of fast and slow muscles of the rabbit. J. Gen. Physiol. 66, 811–821. (doi:10.1085/jgp.66.6.811).

66. Sawicki GS, Sheppard P, Roberts TJ. 2015 Power amplification in an isolated muscle–tendon unit is load dependent. J. Exp. Biol. 218, 3700–3709. (doi:10.1242/jeb.126235).

67. Ellerby DJ, Askew GN. 2007 Modulation of flight muscle power output in budgerigars Melopsittacus undulatus and zebra finches Taeniopygia guttata: in vitro muscle performance. J. Exp. Biol. 210, 3780–3788. (doi:10.1242/jeb.006288).

68. Pollock CM, Shadwick RE. 1994 Allometry of muscle, tendon, and elastic energy storage capacity in mammals. Am. J. Physiol. Regul. Integr. Comp. Physiol. 266, R1022–R1031. (doi:10.1152/ajpregu.1994.266.3.R1022).

69. van der Zee TJ, Wong JD, Kuo AD. 2024 On the rate-limiting dynamics of force development in muscle. J. Exp. Biol. 227, jeb247436. (doi:10.1242/jeb.247436).

70. James RS, Walter I, Seebacher F. 2011 Variation in expression of calcium-handling proteins is associated with inter-individual differences in mechanical performance of rat (Rattus norvegicus) skeletal muscle. J. Exp. Biol. 214, 3542–3548. (doi:10.1242/jeb.058305).

71. Vega D, Arellano CJ. 2026 In situ muscle pre-activation shifts the lateral gastrocnemius muscle–tendon unit to rely on active fascicle lengthening to absorb peak power input. J. Exp. Biol. 229, jeb251324. (doi:10.1242/jeb.251324).

72. Riddell DO, Hildyard JCW, Harron RCM, Taylor-Brown F, Kornegay JN, Wells DJ, Piercy RJ. 2023 Longitudinal assessment of skeletal muscle functional mechanics in the DE50-MD dog model of Duchenne muscular dystrophy. Dis. Model. Mech. 16, dmm050395. (doi:10.1242/dmm.050395).

73. Nguyen A, Leong K, Holt NC. 2025 Does the unusual phenomenon of sustained force circumvent the speed–endurance trade-off in the jaw muscle of the southern alligator lizard (Elgaria multicarinata)? J. Exp. Biol. 228, jeb247979. (doi:10.1242/jeb.247979).

74. Shelley SP, James RS, Eustace SJ, Eyre E, Tallis J. 2022 Effect of stimulation frequency on force, power, and fatigue of isolated mouse extensor digitorum longus muscle. J. Exp. Biol. 225, jeb243285. (doi:10.1242/jeb.243285).

75. James RS, Vanhooydonck B, Tallis JA, Herrel A. 2015 Larger lacertid lizard species produce higher than expected iliotibialis muscle power output: the evolution of muscle contractile mechanics with body size. J. Exp. Biol. 218, 3589–3595. (doi:10.1242/jeb.124974).

76. James RS, Wilson RS. 2008 Explosive jumping: extreme morphological and physiological specializations of Australian rocket frogs (Litoria nasuta). Physiol. Biochem. Zool. 81, 176–185. (doi:10.1086/525290).

77. Alexander RMcN. 1983 Allometry of the leg bones of moas (Dinornithes) and other birds. J. Zool. 200, 215–231. (doi:10.1111/j.1469-7998.1983.tb05785.x).

78. McGowan CP, Skinner J, Biewener AA. 2008 Hind limb scaling of kangaroos and wallabies (superfamily Macropodoidea): implications for hopping performance, safety factor and elastic savings. J. Anat. 212, 153–163. (doi:10.1111/j.1469-7580.2007.00841.x).

79. Alexander RMcN, Jayes AS, Maloiy GMO, Wathuta EM. 1979 Allometry of the limb bones of mammals from shrews (Sorex) to elephant (Loxodonta). J. Zool. 189, 305–314. (doi:10.1111/j.1469-7998.1979.tb03964.x).

80. Clauser CE, McConville JT, Young JW. 1969 Weight, volume, and center of mass of segments of the human body. Aerospace Medical Research Laboratories Technical Report AMRL-TR-69-70 (AD 710 622). Wright-Patterson Air Force Base, OH: Aerospace Medical Research Laboratories, USAF.

81. de Leva P. 1996 Adjustments to Zatsiorsky-Seluyanov’s segment inertia parameters. J. Biomech. 29, 1223–1230. (doi:10.1016/0021-9290(95)00178-6).

82. Grand TI. 1977 Body weight: its relation to tissue composition, segment distribution, and motor function. I. Interspecific comparisons. Am. J. Phys. Anthropol. 47, 211–240. (doi:10.1002/ajpa.1330470204).

83. Isler K, Payne RC, Günther MM, Thorpe SKS, Li Y, Savage R, Crompton RH. 2006 Inertial properties of hominoid limb segments. J. Anat. 209, 201–218. (doi:10.1111/j.1469-7580.2006.00588.x).

84. Schoonaert K, D’Août K, Aerts P. 2007 Morphometrics and inertial properties in the body segments of chimpanzees (Pan troglodytes). J. Anat. 210, 518–531. (doi:10.1111/j.1469-7580.2007.00720.x).

85. Druelle F, Schoonaert K, Aerts P, Nauwelaerts S, Stevens JMG, D’Août K. 2018 Segmental morphometrics of bonobos (Pan paniscus): are they really different from chimpanzees (Pan troglodytes)? J. Anat. 233, 843–853. (doi:10.1111/joa.12894).

