## Supplementary Materials for "Dynamics of Take-off in Bipedal Animals and Robots"

This file includes:

Supplementary Text

Supplementary Figs. 1 to 8

Supplementary Tables 1 to 12

**Initial posture and static equilibrium**

We first determine the horizontal moment arm $D_{c}$ of the upper-body point mass and the initial muscle forces $F_{2,static}$ and $F_{3,static}$ by enforcing static equilibrium at the posture. The position parameters $D$ and $\phi$ were subsequently obtained from $D_{c}$ and the initial geometry.

Static moment balance about the toe, ankle, and knee then gave

$$\left( \frac{1}{2}m_{1}+m_{2}+m_{3}+M_{u} \right)gl_{1}\cos\theta_{1,0}=F_{2,static}l_{1}\sin\left( \theta_{1,0}+\theta_{2,0} \right),$$

(1)

$$\left( \frac{1}{2}m_{2}+m_{3}+M_{u} \right)gl_{2}\cos\theta_{2,0}=F_{3,static}l_{2}\sin\left( \theta_{2,0}+\theta_{3,0} \right)+F_{2,static}r_{2}$$

(2)

$$\frac{1}{2}m_{3}gl_{3}\cos\theta_{3,0}+M_{u}gD_{c}=F_{3,static}r_{3}.$$

(3)

Here, the subscript 0 denotes the initial posture, and $D_{c}$ is the horizontal moment arm of the upper-body point mass about the knee joint. The three static moment-balance equations are solved simultaneously for $F_{2,static}$, $F_{3,static}$, and $D_{c}$.

The upper-body point mass, fixed relative to the thigh, rotates with the thigh about the knee. The knee, hip, and upper-body point mass therefore form a rigid triangle whose shape remain unchanged throughout take-off, as shown in Fig. 1A. Here, $D$ is the distance from the knee joint to the upper-body point mass, $\phi$ is the angle between the thigh segment and the line connecting the knee joint to the upper-body point mass, and $\theta_{4}$ is the prescribed angle at the hip between the thigh and the line from the hip to the upper-body point mass, and is determined from the same sources as the initial segment angles^23,31^. The horizontal projection and geometric relation are

$$D_{c}=D\cos\left( \theta_{3,0}+\phi\right),$$

(4)

$$D=\frac{l_{3}\sin\theta_{4}}{\sin\left( \theta_{4}+\phi\right)},$$

(5)

where Eq. (5) is obtained by applying the law of sines to this triangle. Eqs. (4) and (5) are solved simultaneously for $D$ and $\phi$. The resulting $D$ and $\phi$ are held constant relative to the thigh segment throughout the simulation and therefore do not introduce additional generalized coordinates. The resulting values of $F_{2,static}$ and $F_{3,static}$ are subsequently used to determine the initial activation levels of the two muscle groups.

**Hill force–velocity relationship**

Muscle contraction generates forces that decrease with increasing contraction velocity^40^. This “hyperbolic” force-velocity relationship has been modeled with Hill’s equation^22^:

$$F_{\mathrm{isot}}=F_{0}\frac{1-v/v_{\max}}{1+\left( F_{0}/a \right)\left( v/v_{\max} \right)}.$$

(6)

where $F_{\mathrm{isot}}$ is the isotonic tension that leads to joint rotation, the isometric tension $F_{0}$ defines the tension against which the muscle neither shortens nor lengthens, $v$ is the muscle contraction velocity, and $v_{\max}$ is the maximal velocity reached without load ($F_{\mathrm{isot}}=0$). Parameter $\left( F_{0}/a \right)$ is the inverse of the curvature of the force-velocity relationship^6^.

We transform the variable $v$ into rotational coordinates to apply Hill’s equation to the model. Given the proximity of the tendons connecting the muscles to the joint, the muscle contraction velocity $v$ is approximately equal to the angular velocity of joint rotation $\dot{\theta}$ multiplied by the radius of the bone $r$, $v=r\dot{\theta}$. This relationship arises from the principle that the tangential velocity equals the radius multiplied by the angular velocity for a rotational system. In addition, we multiply the isotonic tension $F_{\mathrm{isot}}$ by the related muscle fiber area $A_{i}$ and multiply $v_{\max}$ by the related muscle fiber length $L_{i}$ so that the units in the equations match each other. The scaling functions for muscle parameters are derived from published empirical data in Supplementary Table 8. Incorporating muscle activation gives the transformed equations of the muscle forces that work for the model.

**Muscle activation**

Muscle requires time to activate during rapid movements^9,26^. Rosario et al.^26^ approximated activation as a linear increase to unity. We follow Rosario et al. and set up muscle activation as a time-dependent, piecewise-linear input. For each muscle group $i$ ($i=2,3$), we define initial activation as the ratio of the force required to maintain static equilibrium at the initial posture to the maximum isometric force,

$$\alpha_{i,0}=\frac{F_{i,static}}{F_{0}A_{i}}, i=2,3.$$

(7)

where $F_{i,static}$ is the muscle force obtained from static moment balance and $F_{0}A_{i}$ is the maximum isometric force of muscle group $i$. Following the onset of take-off, activation increases linearly and is limited to unity,

$$\alpha_{i}\left( t \right)=min\left( \alpha_{i,0}+\frac{t}{T_{\mathrm{act}}},1 \right), i=2,3.$$

(8)

where $T_{\mathrm{act}}$ is the muscle activation timescale that determines the rate of the linear increase in activation. We incorporated Eq. (8) into Eq. (6) to obtain the complete muscle-force equation.

$$F_{2}\left( t \right)=\alpha_{2}\left( t \right)F_{0}A_{2}\frac{1-{r_{2}\left( \dot{\theta}_{1}+\dot{\theta}_{2} \right)}/{v_{\max}L_{2}}}{1+\left( \frac{F_{0}}{a} \right){r_{2}\left( \dot{\theta}_{1}+\dot{\theta}_{2} \right)}/{v_{\max}L_{2}}}.$$

(9)

$$F_{3}\left( t \right)=\alpha_{3}\left( t \right)F_{0}A_{3}\frac{1-{r_{3}\left( \dot{\theta}_{2}+\dot{\theta}_{3} \right)}/{v_{\max}L_{3}}}{1+\left( \frac{F_{0}}{a} \right){r_{3}\left( \dot{\theta}_{2}+\dot{\theta}_{3} \right)}/{v_{\max}L_{3}}}.$$

(10)

Substituting Eqs. (9) and (10) into Eqs. (13) – (15) in Section 4.2 yields the muscle-generated generalized forces for the forward-dynamics simulations.

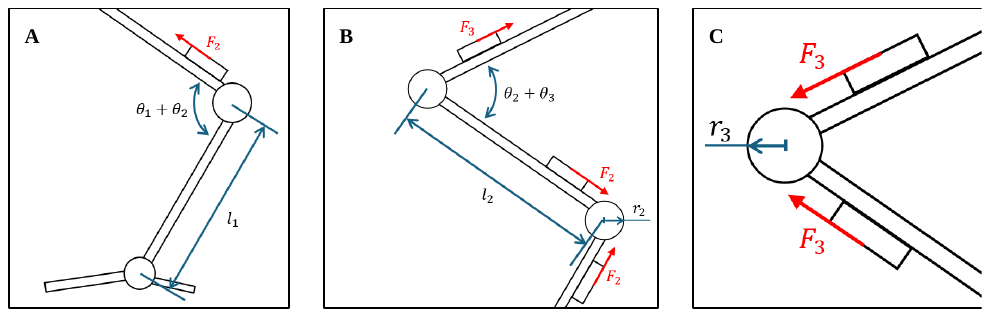

**Supplementary Fig. 1. Free-body diagram of each joint.** (**A**) The torque on toe joint. (**B**) The torque on ankle joint. (**C**) The torque on knee joint.

**A	B**

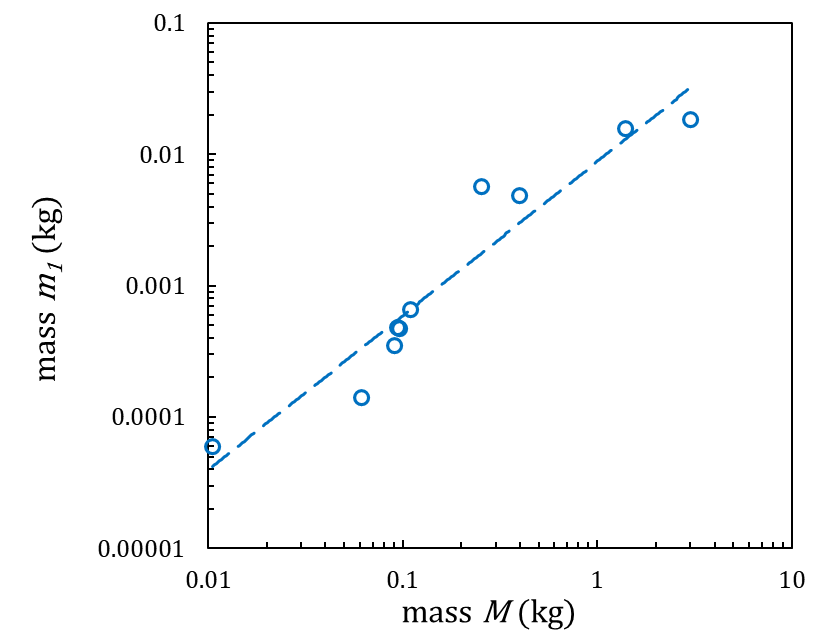

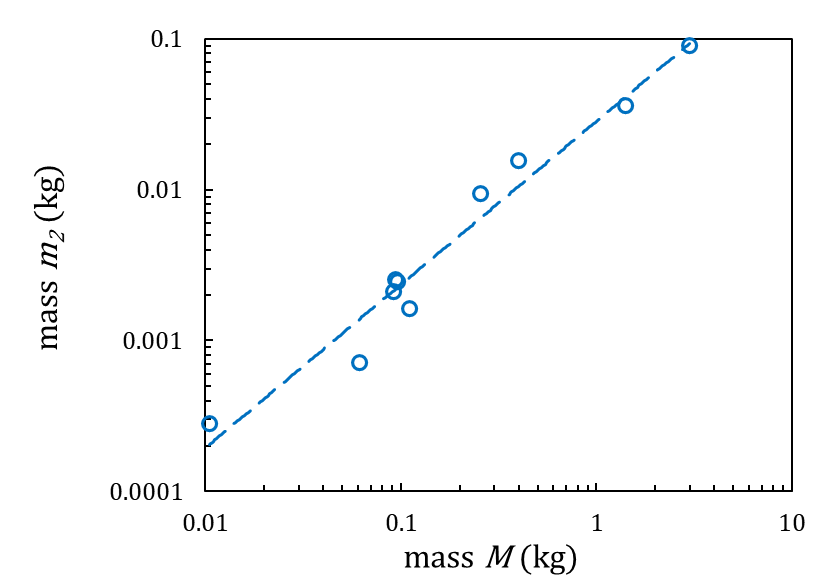

**C**

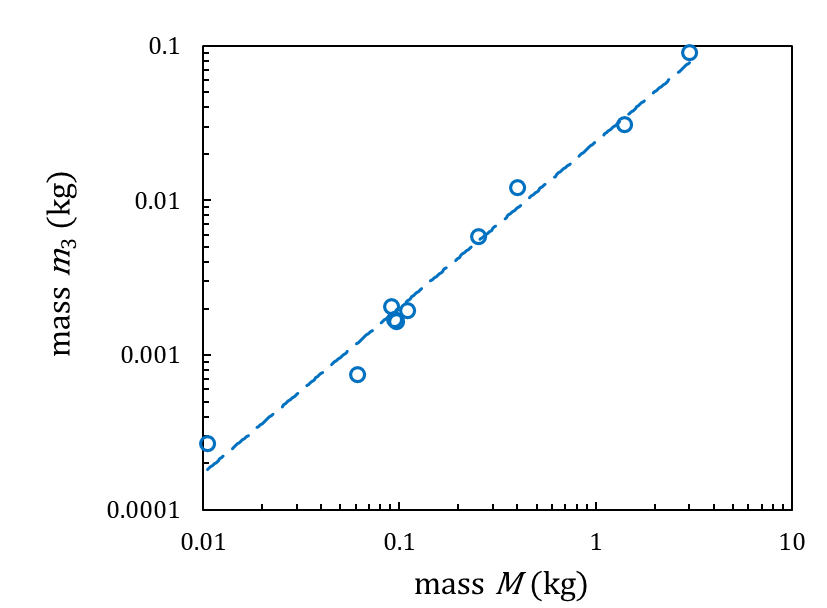

**Supplementary Fig. 2. The relation between mass** $\boldsymbol{M}$ **and the masses of lower limb segments. (A)** mass of tarsus **(B)** mass of shank **(C)** mass of thigh.

**A	B**

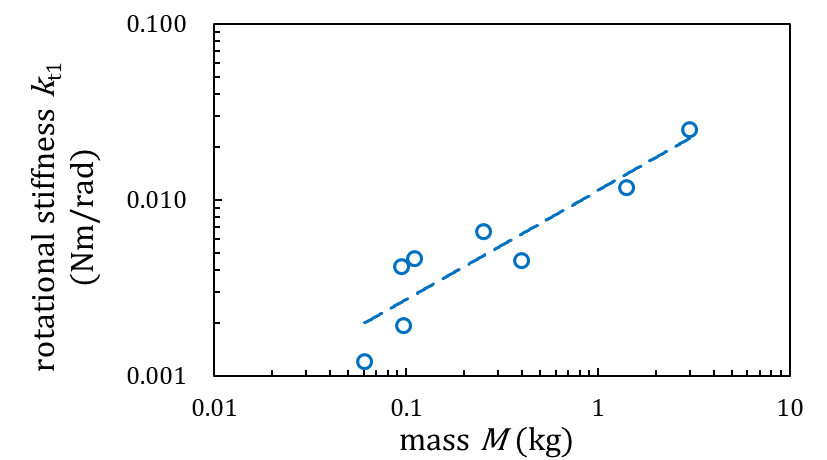

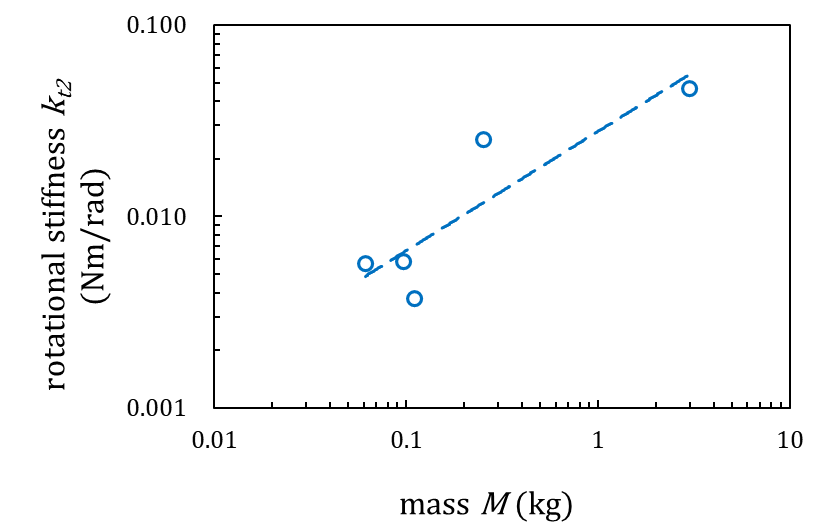

**C**

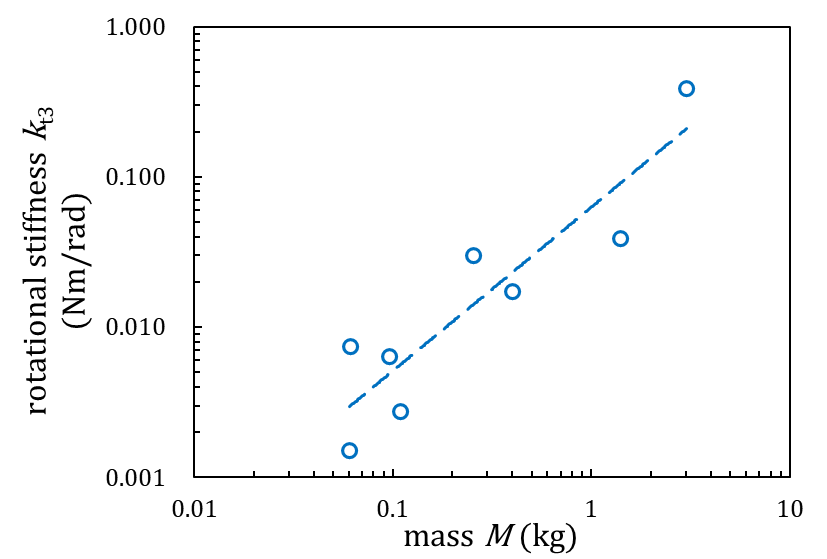

**Supplementary Fig. 3. The relation between mass** $\boldsymbol{M}$ **and rotational stiffness** $\boldsymbol{k}_{\boldsymbol{t}}$ **on each joint. (A)** rotational stiffness at the toe joint **(B)** rotational stiffness at the ankle joint **(C)** rotational stiffness at the knee joint.

**A	B**

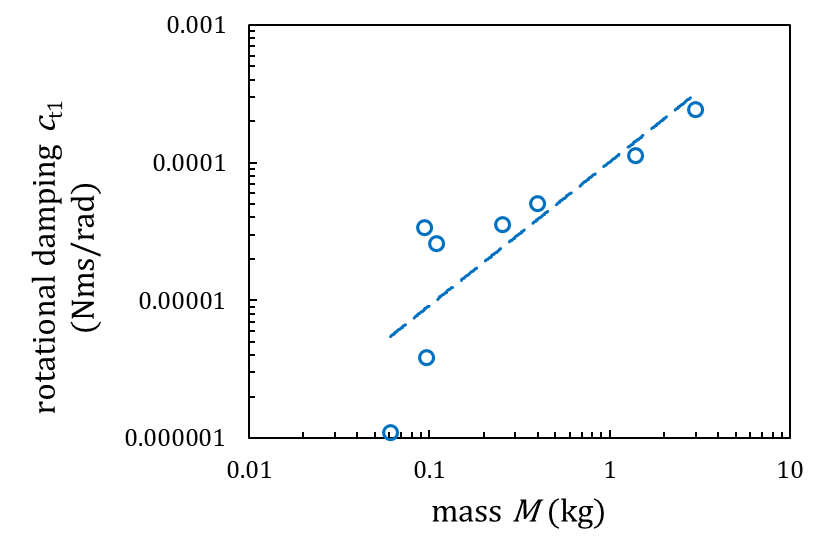

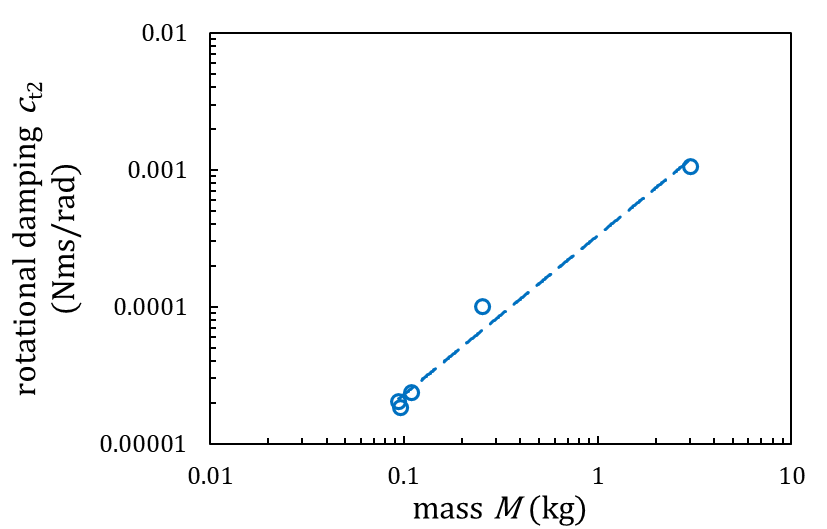

**C**

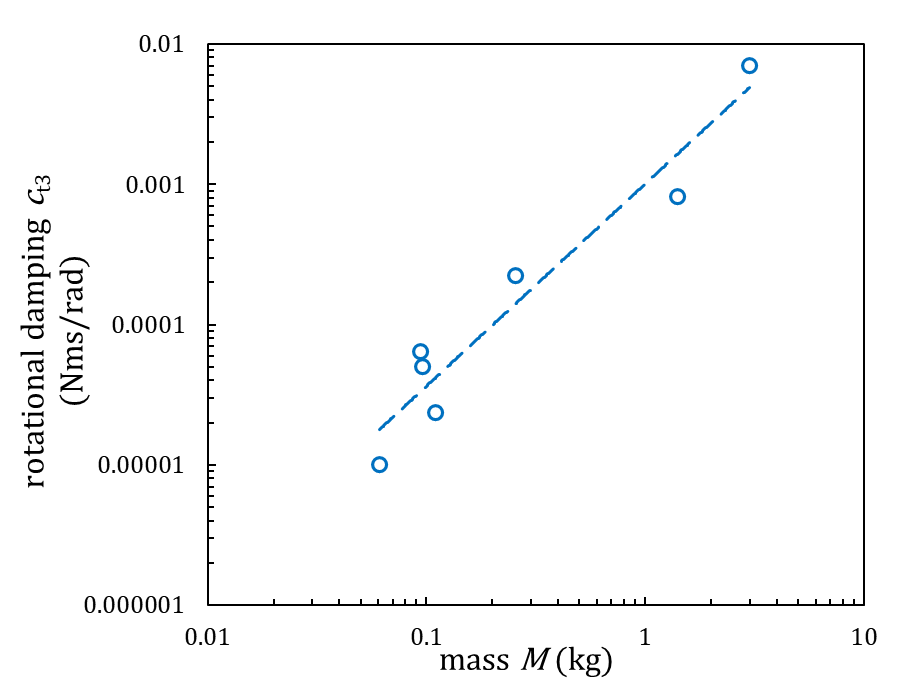

**Supplementary Fig. 4. The relation between mass** $\boldsymbol{M}$ **and rotational damping** $\boldsymbol{c}_{\boldsymbol{t}}$ **on each joint. (A)** rotational damping on the toe joint **(B)** rotational damping on the ankle joint **(C)** rotational damping on the knee joint.

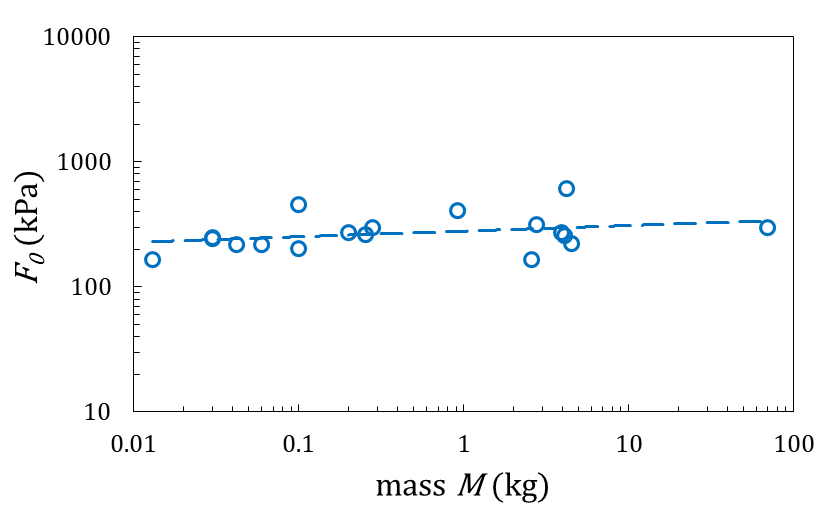

**Supplementary Fig. 5. The relation between mass** $\boldsymbol{M}$ **and maximum isometric tension** $\boldsymbol{F}_{\boldsymbol{0}}$**.**

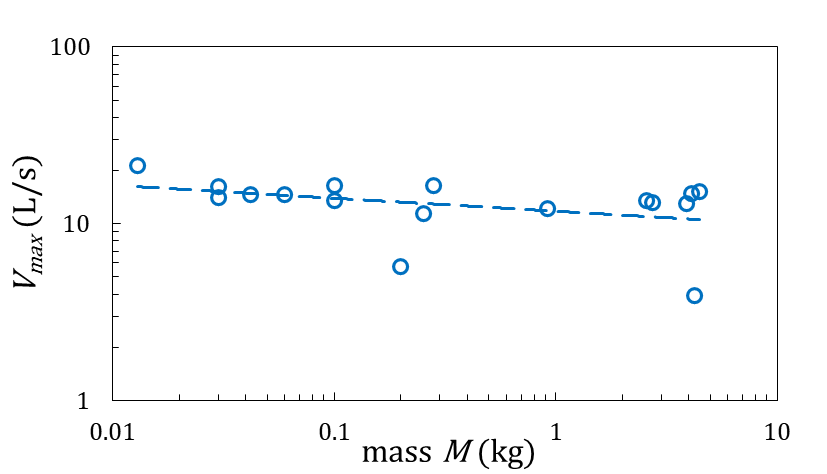

**Supplementary Fig. 6. The relation between mass** $\boldsymbol{M}$ **and maximum muscle contraction velocity** $\boldsymbol{v}_{\boldsymbol{max}}$**.**

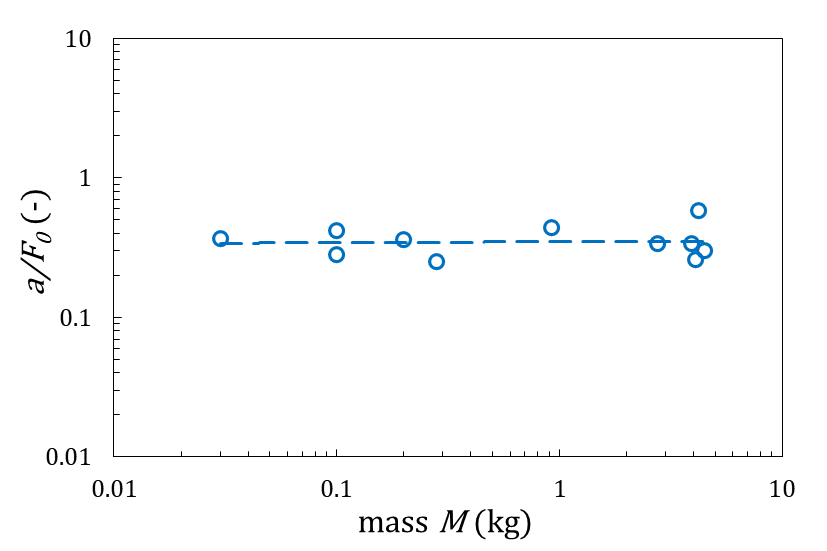

**Supplementary Fig. 7. The relation between mass** $\boldsymbol{M}$ **and curvature of the force-velocity relationship** $\boldsymbol{a}/{\boldsymbol{F}_{\boldsymbol{0}}}$**.**

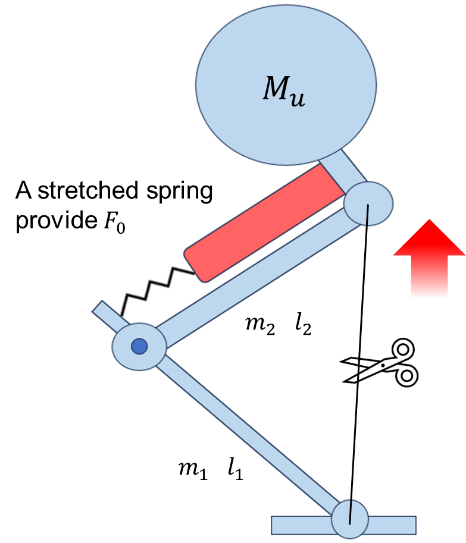

**Supplementary Fig. 8. Schematic of the tabletop mechanism.**

**Supplementary Table 1. List of symbols**

| **Definition** | **symbol** | **unit** |
| --- | --- | --- |
| Distance from the knee joint to the upper-body point mass | $D$ | m |
| Angle between the thigh and the line connecting the knee joint to the upper-body point mass | $\phi$ | radian |
| Passive rotational stiffness | $k_{t}$ | N-m |
| Passive rotational damping coefficient | $c_{t}$ | N-m-s |
| Mass of lower-limb segment i (i = 1, 2, 3) | $m_{i}$ | kg |
| Length of lower-limb segment i (i = 1, 2, 3) | $l_{i}$ | m |
| Half mass of the upper body represented by the upper-body point mass | $M_{u}$ | kg |
| Prescribed hip angle used to define the upper-body geometry | $\theta_{4}$ | radian |
| Angle between the tarsus and the horizontal | $\theta_{1}$ | radian |
| Angle between the shank and the horizontal | $\theta_{2}$ | radian |
| Angle between the thigh and the horizontal | $\theta_{3}$ | radian |
| Horizontal position of the toe | $x_{toe}$ | m |
| Vertical position of the toe | $y_{toe}$ | m |
| Torque on a joint | $\tau$ | N-m |
| Angular displacement | $\Delta\theta$ | degree |
| Young’s modulus of the soft tissues around a joint | $E$ | kPa |
| Cross-sectional area of the soft tissues around a joint | $A$ | m² |
| Effective bone or joint radius | $r$ | m |
| Original length of the soft tissues around a joint | $l_{0}$ | m |
| Body mass | $M$ | kg |
| Rotational stiffness on the toe joint | $k_{t1}$ | N-m |
| Rotational stiffness on the ankle joint | $k_{t2}$ | N-m |
| Rotational stiffness on the knee joint | $k_{t3}$ | N-m |
| Rotational damping coefficient on the toe joint | $c_{t1}$ | N-m-s |
| Rotational damping coefficient on the ankle joint | $c_{t2}$ | N-m-s |
| Rotational damping coefficient on the knee joint | $c_{t3}$ | N-m-s |
| Damping ratio | $\zeta$ | – |
| Moment of inertia | $I$ | kg-m² |
| Critical rotational damping constant | $c_{ct}$ | N-m-s |
| Take-off time | $T$ | ms |
| Maximum ground reaction force | $F$ | N |
| Maximum velocity during take-off | $V_{max}$ | m/s |
| Acceleration due to gravity | $g$ | m/s² |
| Generalized-coordinate vector | $\{q\}$ | – |
| Generalized coordinate i | $q_{i}$ | – |
| Ground-contact constraint Jacobian | $[J_{c}]$ | – |
| Lagrange multipliers associated with horizontal and vertical ground contact | $\lambda_{x}, \lambda_{y}$ | N |
| Vector of ground-contact Lagrange multipliers | $\{\lambda\}$ | N |
| Vector of nonconservative generalized forces | $\{Q\}$ | – |
| Kinetic energy | $KE$ | kg-m/s |
| Potential energy | $V$ | kg-m/s |
| Rayleigh’s dissipation function | $R$ | N-m/s |
| Generalized-coordinate velocity vector | $\{q̇\}$ | – |
| Generalized-coordinate acceleration vector | $\{q̈\}$ | – |
| Velocity of the center of mass of lower-limb segment i | $v_{ci}$ | m/s |
| Moment of inertia of lower-limb segment i about its center of mass | $I_{i}$ | kg-m² |
| Height of the center of mass of lower-limb segment i | $h_{i}$ | m |
| Velocity of the upper-body point mass | $v_{cu}$ | m/s |
| Height of the upper-body point mass | $h_{u}$ | m |
| Inertia matrix | $[I]$ | – |
| Coriolis, centrifugal, and damping matrix | $[C]$ | – |
| Stiffness matrix | $[G_{s}]$ | – |
| Gravitational matrix | $[G_{g}]$ | – |
| Ground reaction force in the x-direction | $F_{x}$ | N |
| Ground reaction force in the y-direction | $F_{y}$ | N |
| Torque on the toe joint | $\tau_{1}$ | N-m |
| Torque on the ankle joint | $\tau_{2}$ | N-m |
| Torque on the knee joint | $\tau_{3}$ | N-m |
| Muscle force from the calf | $F_{2}$ | N |
| Muscle force from the thigh | $F_{3}$ | N |
| Radius of the ankle joint | $r_{2}$ | m |
| Radius of the knee joint | $r_{3}$ | m |
| Static muscle force from the calf | $F_{2,static}$ | N |
| Static muscle force from the thigh | $F_{3,static}$ | N |
| Horizontal moment arm of the upper-body point mass about the knee joint | $D_{c}$ | m |
| Initial angle of lower-limb segment i | $\theta_{i,0}$ | radian |
| Isotonic tension | $F_{isot}$ | N/m² |
| Maximum isometric tension | $F_{0}$ | N/m² |
| Muscle contraction velocity | $v$ | L/s |
| Maximum muscle contraction velocity | $v_{max}$ | L/s |
| The inverse of the curvature of the force-velocity relationship | $F_{0}/a$ | – |
| Muscle fiber area related to muscle force Fᵢ (i = 2, 3) | $A_{i}$ | m² |
| Muscle fiber length related to muscle force Fᵢ (i = 2, 3) | $L_{i}$ | m |
| Initial activation of muscle group i | $\alpha_{i,0}$ | – |
| Time-dependent activation of muscle group i | $\alpha_{i}(t)$ | – |
| Muscle activation timescale | $T_{act}$ | s |
| Stretching force in the soft tissues around a joint | $f$ | N |
| Spring constant of the soft tissues around a joint | $k$ | N/m |
| Logarithmic decrement | $\delta$ | – |
| Amplitudes of two adjacent free vibrations | $x_{1}, x_{2}$ | degree |
| Initial generalized-coordinate vector | $\{q_{0}\}$ | – |
| Radius of the tibia shaft | $r_{2'}$ | m |
| Radius of the femur shaft | $r_{3'}$ | m |

**Supplementary Table 2. Take-off time across bipedal species (n=38)**

| **species** | **mass**  **(kg)** | **time (ms)** | **sources** |
| --- | --- | --- | --- |
| Hummingbird (*Selasphorus rufus*) | 0.003 | 71 | (44) |
| Zebra Finch (*Taeniopygia guttata*) | 0.015 | 28 | (45) |
| Sparrow (*Passer montanus*) | 0.024 | 50 | ^*^YouTube,  Smarter Every Day 2 |
| Diamond Dove (*Geopelia cuneata*) | 0.051 | 39 | (45) |
| White-headed Buffalo-Weaver  (*Dinemellia dinemelli*) | 0.064 | 120 | Experiment at Zoo Atlanta |
| Superb Starling (*Lamprotornis superbus*) | 0.065 | 128 | Experiment at Zoo Atlanta |
| Wood Hoopoe (*Phoeniculus purpureus*) | 0.075 | 48 | Experiment at Zoo Atlanta |
| European Starling (*Sturnus vulgaris*) | 0.077 | 108 | (46) |
| Quail (*Coturnix coturnix*) | 0.172 | 71 | (46) |
| Golden-backed Weaverbird  (*Ploceus jacksoni*) | 0.220 | 57 | Experiment at Taipei Zoo |
| Rose-breasted Cockatoo  (*Eolophus roseicapilla*) | 0.310 | 138 | Experiment at Taipei Zoo |
| White-cheeked Turaco  (*Tauraco leucotis*) | 0.323 | 130 | Experiment at Zoo Atlanta |
| Laughing Kookaburra  (*Dacelo novaeguineae*) | 0.348 | 200 | Experiment at Zoo Atlanta |
| Speckled Pigeon  (*Columba guinea*) | 0.352 | 150 | Experiment at Zoo Atlanta |
| Orange-crested Cockatoo  (*Cacatua sulphurea*) | 0.400 | 136 | Experiment at Taipei Zoo |
| Sun Conure (*Aratinga solstitialis*) | 0.400 | 138 | Experiment at Taipei Zoo |
| Blue-fronted Amazon (*Amazona aestiva*) | 0.420 | 122 | Experiment at Taipei Zoo |
| Pied Imperial Pigeon  (*Ducula bicolor*) | 0.440 | 88 | Experiment at Taipei Zoo |
| Muscovy Duck (*Cairina moschata*) | 0.450 | 138 | Experiment at Taipei Zoo |
| Barn Owl (*Tyto alba*) | 0.470 | 140 | Experiment at Zoo Atlanta |
| African Grey Parrot (*Psittacus erithacus*) | 0.500 | 99 | Experiment at Taipei Zoo |
| Senegal Parrot (*Poicephalus senegalus*) | 0.500 | 121 | Experiment at Taipei Zoo |
| Bald Eagle (*Haliaeetus leucocephalus*) | 0.500 | 119 | Experiment at Taipei Zoo |
| Nicobar Pigeon (*Caloenas nicobarica*) | 0.500 | 130 | Experiment at Taipei Zoo |
| White-headed Marmoset  (*Callithrix geoffroyi*) | 0.549 | 125 | (47) |
| Yellow-naped Amazon  (*Amazona auropalliata*) | 0.580 | 129 | Experiment at Taipei Zoo |
| Palm Cockatoo (*Probosciger aterrimus*) | 0.750 | 120 | Experiment at Taipei Zoo |
| Red Avadavat (*Amandava amandava*) | 0.800 | 100 | Experiment at Taipei Zoo |
| Verreaux's Eagle-Owl (*Bubo lacteus*) | 1.18 | 184 | Experiment at Zoo Atlanta |
| Guinea Fowl (*Numida meleagris*) | 1.42 | 130 | (23) |
| Brown Wood Owl  (*Strix leptogrammica*) | 1.50 | 135 | Experiment at Taipei Zoo |
| Philippine Eagle  (*Pithecophaga jefferyi*) | 1.60 | 125 | Experiment at Taipei Zoo |
| Hooded Vulture  (*Necrosyrtes monachus*) | 1.72 | 142 | Experiment at Zoo Atlanta |
| Tawny Fish Owl (*Ketupa flavipes*) | 2.60 | 143 | Experiment at Taipei Zoo |
| Harpy Eagle (*Harpia harpyja*) | 2.20 | 128 | Experiment at Taipei Zoo |
| Rock Wallaby (*Petrogale xanthopus*) | 5.50 | 71 | (48) |
| Indri (*Indri indri*) | 6.50 | 210 | (49) |
| Human02 (*Homo sapiens*) | 90.0 | 294 | ^*^YouTube, LocomoterLabTCU |

Smarter Every Day 2. (2018, Aug 8). Bird Taking Off at 20,000 fps (213milliseconds) Raw Video - Smarter Every Day 197. [Video]. YouTube.

<https://youtu.be/EjVvdkKBh2w?si=MbE8uIo6fllT3eNL>

LocomoterLabTCU. (2012, Sep 29). Slow-motion video of vertical jump with synchronized vertical force data. [Video]. YouTube.

<https://youtu.be/qN3apht8zRs?si=vDpi4An6X-Vt7D1j>

**Supplementary Table 3. Maximum ground reaction force across bipedal species (n=11)**

| **species** | **mass (kg)** | **force (N)** | **sources** |
| --- | --- | --- | --- |
| Hummingbird (*Selasphorus rufus*) | 0.003 | 0.04 | (44) |
| Zebra Finch (*Taeniopygia guttata*) | 0.015 | 0.74 | (45) |
| Diamond Dove (*Geopelia cuneata*) | 0.051 | 1.36 | (45) |
| Leopard Frog  (*Lithobates sphenocephalus*) | 0.055 | 2.63 | (5) |
| Starling (*Sturnus vulgaris*) | 0.077 | 3.29 | (46) |
| Quail (*Coturnix coturnix*) | 0.172 | 13.9 | (46) |
| White-headed Marmoset  (*Callithrix geoffroyi*) | 0.549 | 26.0 | (47) |
| Guinea fowl (*Numida meleagris*) | 1.42 | 78.0 | (23) |
| Yellow-footed rock wallaby  (*Petrogale xanthopus*) | 5.50 | 451 | (48) |
| Bonobo (*Pan paniscus*) | 34 | 867 | (50) |
| Human (*Homo sapiens*) | 75.5 | 1.85$\times$10^3^ | (29) |

**Supplementary Table 4. Maximum velocity across bipedal species (n=13)**

| **species** | **mass (kg)** | **velocity (m/s)** | **sources** |
| --- | --- | --- | --- |
| Hummingbirds (*Selasphorus rufus*) | 0.003 | 0.70 | (44) |
| Zebra Finch (*Taeniopygia guttata*) | 0.015 | 1.74 | (45) |
| Kangaroo rat (*Dipodomys merriami*) | 0.035 | 1.76 | (51) |
| Blue-breasted quail (*Coturnix chinensis*) | 0.044 | 0.78 | (52) |
| Diamond Dove (*Geopelia cuneata*) | 0.051 | 1.29 | (45) |
| Starling (*Sturnus vulgaris*) | 0.079 | 2.21 | (53) |
| Kangaroo rat (*Dipodomys spectabilis*) | 0.099 | 2.12 | (51) |
| Quail (*Coturnix coturnix*) | 0.172 | 3.79 | (46) |
| White-headed Marmoset  (*Callithrix geoffroyi*) | 0.549 | 3.39 | (47) |
| Guinea fowl (*Numida meleagris*) | 1.42 | 3.60 | (23) |
| Yellow-footed rock wallaby  (*Petrogale xanthopus*) | 5.50 | 5.23 | (48) |
| Bonobo (*Pan paniscus*) | 34 | 3.71 | (50) |
| Human (*Homo sapiens*) | 75.5 | 2.69 | (29) |

**Supplementary Table 5. The lower limb segment masses of Aves (n=11)**

| **species** | **mass (kg)** | $\boldsymbol{m}_{\boldsymbol{1}}$ **(g)** | $\boldsymbol{m}_{\boldsymbol{2}}$ **(g)** | $\boldsymbol{m}_{\boldsymbol{3}}$ **(g)** |
| --- | --- | --- | --- | --- |
| Sparrow (*Passer montanus*) | 0.011 | 0.06 | 0.28 | 0.27 |
| Muller’s barbet 01  (*Psilopogon nuchalis*) | 0.061 | 0.35 | 1.75 | 1.31 |
| Black bulbul  (*Hypsipetes leucocephalus*) | 0.061 | 0.14 | 0.72 | 0.75 |
| Grey treepie 01  (*Dendrocitta formosae*) | 0.091 | 0.35 | 2.11 | 2.05 |
| Muller’s Barbet 02  (*Psilopogon nuchalis*) | 0.094 | 0.48 | 2.53 | 1.71 |
| Grey treepie 02  (*Dendrocitta formosae*) | 0.096 | 0.47 | 2.47 | 1.65 |
| Spotted dove  (*Spilopelia chinensis*) | 0.11 | 0.66 | 1.62 | 1.94 |
| Malaysian night heron  (*Gorsachius melanolophus*) | 0.254 | 5.65 | 9.43 | 5.86 |
| Cattle egret  (*Bubulcus coromandus*) | 0.4 | 4.83 | 15.59 | 12.08 |
| Sacred ibis  (*Threskiornis aethiopicus*) | 1.4 | 15.72 | 36.28 | 30.74 |
| Chicken  (*Gallus gallus domesticus*) | 3 | 18.3 | 90 | 90 |

**Supplementary Table 6. The rotational stiffness of different joints (n=9)**

| **species** | **mass (kg)** | **joint** | $\boldsymbol{k}_{\boldsymbol{t}}$ **(Nm/rad)** |
| --- | --- | --- | --- |
| Muller’s barbet 01  (*Psilopogon nuchalis*) | 0.061 | ankle | $5.70\times{10}^{-3}$ |
|  |  | knee | $7.40\times{10}^{-3}$ |
| Grey treepie  (*Dendrocitta formosae*) | 0.096 | toe | $1.92\times{10}^{-3}$ |
|  |  | ankle | $5.80\times{10}^{-3}$ |
|  |  | knee | $6.40\times{10}^{-3}$ |
| Spotted dove  (*Spilopelia chinensis*) | 0.110 | toe | $4.64\times{10}^{-3}$ |
|  |  | ankle | $3.74\times{10}^{-3}$ |
|  |  | knee | $2.74\times{10}^{-3}$ |
| Cattle egret  (*Bubulcus coromandus*) | 0.400 | toe | $4.50\times{10}^{-3}$ |
|  |  | knee | $17.3\times{10}^{-3}$ |
| Chicken  (*Gallus gallus domesticus*) | 3 | toe | $24.9\times{10}^{-3}$ |
|  |  | ankle | $46.8\times{10}^{-3}$ |
|  |  | knee | $389\times{10}^{-3}$ |
| Sacred ibis  (*Threskiornis aethiopicus*) | 1.4 | toe | $11.7\times{10}^{-3}$ |
|  |  | knee | $38.9\times{10}^{-3}$ |
| Black bulbul  (*Hypsipetes leucocephalus*) | 0.061 | toe | $1.20\times{10}^{-3}$ |
|  |  | knee | $1.50\times{10}^{-3}$ |
| Malaysian night heron  (*Gorsachius melanolophus*) | 0.254 | toe | $6.60\times{10}^{-3}$ |
|  |  | ankle | $2.52\times{10}^{-3}$ |
|  |  | knee | $2.98\times{10}^{-3}$ |
| Muller’s barbet 02  (*Psilopogon nuchalis*) | 0.094 | toe | $4.18\times{10}^{-3}$ |

**Supplementary Table 7. The rotational damping of different joints (n=8)**

| **species** | **mass (kg)** | **joint** | $\boldsymbol{c}_{\boldsymbol{t}}$ **(Nms/rad)** |
| --- | --- | --- | --- |
| Muller’s barbet  (*Psilopogon nuchalis*) | 0.094 | toe | $3.40\times{10}^{-5}$ |
|  |  | ankle | $2.04\times{10}^{-5}$ |
|  |  | knee | $6.46\times{10}^{-5}$ |
| Grey treepie  (*Dendrocitta formosae*) | 0.096 | toe | $3.82\times{10}^{-6}$ |
|  |  | ankle | $1.84\times{10}^{-5}$ |
|  |  | knee | $5.04\times{10}^{-5}$ |
| Spotted dove  (*Spilopelia chinensis*) | 0.110 | toe | $2.60\times{10}^{-5}$ |
|  |  | ankle | $2.38\times{10}^{-5}$ |
|  |  | knee | $2.35\times{10}^{-5}$ |
| Cattle egret  (*Bubulcus coromandus*) | 0.400 | toe | $5.01\times{10}^{-5}$ |
| Chicken  (*Gallus gallus domesticus*) | 3 | toe | $2.43\times{10}^{-4}$ |
|  |  | ankle | $1.06\times{10}^{-3}$ |
|  |  | knee | $7.07\times{10}^{-3}$ |
| Sacred ibis  (*Threskiornis aethiopicus*) | 1.4 | toe | $1.14\times{10}^{-4}$ |
|  |  | knee | $8.11\times{10}^{-4}$ |
| Black bulbul  (*Hypsipetes leucocephalus*) | 0.061 | toe | $1.10\times{10}^{-6}$ |
|  |  | knee | $9.98\times{10}^{-6}$ |
| Malaysian night heron  (*Gorsachius melanolophus*) | 0.254 | toe | $3.53\times{10}^{-5}$ |
|  |  | ankle | $1.01\times{10}^{-4}$ |
|  |  | knee | $2.22\times{10}^{-4}$ |

**Supplementary Table 8.** **The common scaling function for all bipedal animals in the compound pendulum model**

|  | scaling function | n | unit | sources |
| --- | --- | --- | --- | --- |
| rotational stiffness on toe joint | $k_{t1}=0.032M^{0.52}$ | 9 | Nm | experiments |
| rotational damping on toe joint | $c_{t1}={3.86\times{10}^{-4}M}^{1.23}$ | 8 | Nms | experiments |
| rotational stiffness on ankle | $k_{t2}=0.029M^{0.63}$ | 9 | Nm | experiments |
| rotational damping on ankle | $c_{t2}={4.02\times{10}^{-4}M}^{1.57}$ | 8 | Nms | experiments |
| rotational stiffness on knee | $k_{t3}=0.059M^{1.09}$ | 9 | Nm | experiments |
| rotational damping on knee | $c_{t3}={9.71\times{10}^{-4}M}^{1.44}$ | 8 | Nms | experiments |
| maximum isometric tension | $F_{0}={262\times{10}^{3}M}^{0.03}$ | 18 | N/ m^2^ | (54-67) |
| gastrocnemius muscle fiber area | $A={3.04\times{10}^{-4}M}^{0.77}$ | 26 | m^2^ | (68) |
| maximum muscle contraction velocity | $v_{max}={12.42M}^{-0.025}$ | 17 | L/s | (54-67) |
| gastrocnemius muscle fiber length | $L=10.9\times{10}^{-3}M^{0.21}$ | 26 | m | (68) |
| the curvature of the  force-velocity relationship | $a/F_{0}={0.347M}^{-0.004}$ | 11 | $-$ | (54-67) |
| time until maximum muscle activation | $T_{act}={0.182M}^{0.261}$ | 9 | s | (69-76) |

**Supplementary Table 9. The skeletal scaling functions for three different categories in the compound pendulum model**

|  |  | scaling function | n | unit | sources |
| --- | --- | --- | --- | --- | --- |
| Aves | tarsometatarsus | $l_{1}=0.075M^{0.34}$ | 13 | m | (77) |
|  | tibiotarsus | $l_{2}=0.112M^{0.37}$ | 14 |  |  |
|  | femur | $l_{3}=0.067M^{0.37}$ | 14 |  |  |
|  | radius of tibia shaft | ${r_{2}}^{'}=2.69\times{10}^{-3}M^{0.39}$ | 14 |  |  |
|  | radius of femur shaft | ${r_{3}}^{'}=3.19\times{10}^{-3}M^{0.42}$ | 15 |  |  |
| Macropodidae | metatarsal | $l_{1}=0.04M^{0.37}$ | 21 |  | (78) |
|  | tibia | $l_{2}=0.092M^{0.42}$ |  |  |  |
|  | femur | $l_{3}=0.08M^{0.32}$ |  |  |  |
|  | radius of tibia shaft | ${r_{2}}^{'}=2.47\times{10}^{-3}M^{0.4}$ |  |  |  |
|  | radius of femur shaft | ${r_{3}}^{'}=3.18\times{10}^{-3}M^{0.34}$ |  |  |  |
| Primates | metatarsal | $l_{1}=0.025M^{0.31}$ | 6 |  | (79) |
|  | tibia | $l_{2}=0.092M^{0.32}$ |  |  |  |
|  | femur | $l_{3}=0.094M^{0.36}$ |  |  |  |
|  | radius of tibia shaft | ${r_{2}}^{'}=4.8\times{10}^{-3}M^{0.36}$ |  |  |  |
|  | radius of femur shaft | ${r_{3}}^{'}=5.2\times{10}^{-3}M^{0.36}$ |  |  |  |

**Supplementary Table 10. The lower limb segment mass scaling functions for three different categories in the compound pendulum model**

|  |  | scaling function | n | unit | sources |
| --- | --- | --- | --- | --- | --- |
| Aves | tarsometatarsus | $m_{1}=0.013M^{1.33}$ | 11 | kg | experiments |
|  | shank | $m_{2}=0.032M^{1.13}$ |  |  |  |
|  | thigh | $m_{3}=0.025M^{1.1}$ |  |  |  |
| Macropodidae | metatarus | $m_{1}=0.123M$ | 1 |  | (30) |
|  | shank | $m_{2}=0.045M$ |  |  |  |
|  | thigh | $m_{3}=0.011M$ |  |  |  |
| Primates | foot | $m_{1}=0.017M^{0.96}$ | 27 |  | (80-85) |
|  | shank | $m_{2}=0.022M^{1.05}$ |  |  |  |
|  | thigh | $m_{3}=0.04M^{1.09}$ |  |  |  |

**Supplementary Table 11. Simulation for take-off time, maximum velocity, and maximum ground reaction force**

| **species** | **mass (kg)** | $\boldsymbol{T}$  **(ms)** | $\boldsymbol{V}_{\boldsymbol{max}}$  **(m/s)** | $\boldsymbol{F}$  **(N)** |
| --- | --- | --- | --- | --- |
| Hummingbird (*Selasphorus rufus*) | 0.003 | 25.8 | 0.48 | 0.14 |
| Zebra Finch (*Taeniopygia guttata*) | 0.015 | 40.3 | 0.71 | 0.60 |
| Sparrow (*Passer montanus*) | 0.024 | 45.7 | 0.77 | 0.90 |
| Diamond Dove (*Geopelia cuneata*) | 0.051 | 56.4 | 0.87 | 1.78 |
| White-headed Buffalo-Weaver  (*Dinemellia dinemelli*) | 0.064 | 60.0 | 0.90 | 2.19 |
| Superb Starling (*Lamprotornis superbus*) | 0.065 | 60.2 | 0.91 | 2.22 |
| Wood Hoopoe (*Phoeniculus purpureus*) | 0.075 | 62.7 | 0.92 | 2.52 |
| European Starling (*Sturnus vulgaris*) | 0.077 | 63.1 | 0.93 | 2.58 |
| Quail (*Coturnix coturnix*) | 0.172 | 78.6 | 1.03 | 5.31 |
| Golden-backed Weaverbird  (*Ploceus jacksoni*) | 0.220 | 84.0 | 1.06 | 6.61 |
| Rose-breasted Cockatoo  (*Eolophus roseicapilla*) | 0.310 | 92.2 | 1.10 | 8.98 |
| White-cheeked Turaco (*Tauraco leucotis*) | 0.323 | 93.2 | 1.10 | 9.32 |
| Laughing Kookaburra  (*Dacelo novaeguineae*) | 0.348 | 95.1 | 1.11 | 9.96 |
| Speckled Pigeon (*Columba guinea*) | 0.352 | 95.4 | 1.12 | 10.06 |
| Orange-crested Cockatoo  (*Cacatua sulphurea*) | 0.400 | 98.7 | 1.13 | 11.28 |
| Sun Conure (*Aratinga solstitialis*) | 0.400 | 98.7 | 1.13 | 11.28 |
| Blue-fronted Amazon (*Amazona aestiva*) | 0.420 | 100.0 | 1.14 | 11.78 |
| Pied Imperial Pigeon  (*Ducula bicolor*) | 0.440 | 101.3 | 1.14 | 12.28 |
| Muscovy Duck (*Cairina moschata*) | 0.450 | 101.9 | 1.14 | 12.53 |
| Barn Owl (*Tyto alba*) | 0.470 | 103.1 | 1.15 | 13.03 |
| African Grey Parrot (*Psittacus erithacus*) | 0.500 | 104.9 | 1.16 | 13.77 |
| Senegal Parrot (*Poicephalus senegalus*) | 0.500 | 104.9 | 1.16 | 13.77 |
| Bald Eagle (*Haliaeetus leucocephalus*) | 0.500 | 104.9 | 1.16 | 13.77 |
| Nicobar Pigeon (*Caloenas nicobarica*) | 0.500 | 104.9 | 1.16 | 13.77 |
| White-headed Marmoset  (*Callithrix geoffroyi*) | 0.549 | 110.9 | 0.73 | 11.73 |
| Yellow-naped Amazon  (*Amazona auropalliata*) | 0.580 | 109.2 | 1.17 | 15.72 |
| Palm Cockatoo (*Probosciger aterrimus*) | 0.750 | 117.0 | 1.21 | 19.78 |
| Red Avadavat (*Amandava amandava*) | 0.850 | 119.1 | 1.21 | 20.95 |
| Verreaux's Eagle-Owl (*Bubo lacteus*) | 1.18 | 132.3 | 1.26 | 29.66 |
| Guinea Fowl (*Numida meleagris*) | 1.42 | 139.1 | 1.28 | 35.00 |
| Brown Wood Owl (*Strix leptogrammica*) | 1.50 | 141.2 | 1.29 | 36.76 |
| Philippine Eagle (*Pithecophaga jefferyi*) | 1.60 | 143.7 | 1.30 | 38.95 |
| Hooded Vulture (*Necrosyrtes monachus*) | 1.72 | 146.6 | 1.30 | 41.55 |
| Tawny Fish Owl (*Ketupa flavipes*) | 2.20 | 156.7 | 1.33 | 51.81 |
| Harpy Eagle (*Harpia harpyja*) | 2.20 | 156.7 | 1.33 | 51.81 |
| Rock Wallaby (*Petrogale xanthopus*) | 5.50 | 175.8 | 1.02 | 111.37 |
| Indri (*Indri indri*) | 6.50 | 214.1 | 0.91 | 111.53 |
| Human (*Homo sapiens*) | 90.0 | 467.5 | 1.10 | 1268.98 |
| *Ornitholestes* | 16.5 | 273.0 | 1.59 | 318.769 |
| *Saurornitholestes* | 22.5 | 297.7 | 1.63 | 422.511 |
| *Oviraptor* | 58 | 388.7 | 1.76 | 1002.661 |
| *Ornithomimus* | 155 | 514.9 | 1.90 | 2475.552 |
| *Dromiceiomimus* | 160 | 519.7 | 1.91 | 2549.181 |
| *Anserimimus* | 170 | 528.8 | 1.92 | 2695.962 |
| *Strutiomimus* | 175 | 533.3 | 1.92 | 2769.126 |
| *Elaphrosaurus* | 245 | 588.0 | 1.97 | 3780.348 |
| *Dilophosaurus* | 325 | 638.7 | 2.02 | 4913.173 |
| *Gallimimus* | 490 | 721.3 | 2.08 | 7198.022 |
| *Allosaurus fragilis* | 1620 | 1031.2 | 2.29 | 22051.235 |
| *Tarbosaurus* | 1650 | 1037.0 | 2.29 | 22435.302 |
| *Albertosaurus* | 1685 | 1043.5 | 2.29 | 22882.939 |
| *Sinraptor dongi* | 1700 | 1046.3 | 2.29 | 23074.639 |
| *Daspletosaurus* | 2700 | 1203.7 | 2.38 | 35693.311 |
| *Tyrannosaurus* | 6300 | 1562.0 | 2.54 | 79687.608 |

**Supplementary Table 12. Take-off time for tabletop mechanisms**

| Mass (kg) | First trial  (ms) | Second trial  (ms) | Third trial  (ms) | Average  (ms) | Standard deviation (ms) |
| --- | --- | --- | --- | --- | --- |
| 0.025 | 51 | 50 | 49 | 50 | 1.0 |
| 0.25 | 116 | 122 | 128 | 122 | 6.0 |
| 2.5 | 142 | 153 | 167 | 154 | 12.5 |
